# Tissue geometry patterns epidermal stem cell fate in murine interfollicular skin

**DOI:** 10.64898/2026.11.01.763461

**Authors:** Shicong Xie, Jonathan Karin, Vanessa Lopez-Pajares, Katie Cockburn, Mor Nitzan, Jan M. Skotheim

## Abstract

How tissues maintain spatial patterns of stem cell renewal and differentiation despite continuous cellular turnover remains poorly understood. Here, we show that microscopic undulations of the dermal-epidermal junction (DEJ) are associated with distinct stem cell behaviors in the mouse epidermis. Using longitudinal intravital imaging and quantitative analysis of cell morphology and lineage dynamics, we find that local DEJ curvature predicts the subsequent fate of daughter cells up to three days before mother cell division. Cells residing in concave valleys preferentially produce proliferating descendants, whereas convex ridges are associated with increased differentiation. These spatial biases are reflected in the expression of stem- and differentiation-associated markers and accumulate into differences in long-term clonal expansion. Remarkably, DEJ geometry remains stable for weeks, persisting across multiple generations of epidermal cells. Moreover, human epidermal equivalents grown on devitalized dermis recapitulate the preferential retention of basal layer cells in valleys, indicating that living dermal cells are not required for this spatial organization. Our findings suggest that stable tissue architecture can pattern stem cell fate probabilities across generations, contributing to functional stem cell heterogeneity.

## Introduction

Tissue-resident stem and progenitor cells must continually balance self-renewal with terminal differentiation. This balance maintains tissue architecture despite the persistent loss of mature cells, and its disruption can lead to degeneration, impaired repair, or tumorigenesis. Although biochemical signals regulating stem cell fate have been studied extensively, cells also encounter spatially patterned physical cues imposed by the tissue’s architecture.^1^ Whether such architectural features are merely the consequences of ongoing cell behaviors or instead provide stable spatial information that organizes fate decisions remains poorly understood.

The mammalian interfollicular epidermis provides an accessible *in vivo* system in which to investigate how the physical microenvironment shapes cell fate (**Figure 1A**). During homeostasis, proliferative cells within the basal layer divide to replace biomass lost through shedding at the skin surface. Cells committed to differentiation delaminate apically from the basal layer, enter the suprabasal layers, and progress through a terminal differentiation program (**Figure 1B**). Despite occupying a common compartment, molecular profiling, lineage tracing, and label retention experiments show that basal layer cells exhibit substantial heterogeneity in their transcriptional states, cell cycle behaviors, and division rates.^2,3^ This heterogeneity has raised questions about whether the basal layer comprises functionally distinct progenitor populations or a single population capable of adopting different cellular fates. In the two-progenitor model, the basal layer contains distinct populations of slow- and fast-cycling progenitors with different capacities for long-term self-renewal.^4–7^ Yet, the molecular origin of the signals directing these two populations is unclear. In contrast, the equipotent-progenitor model proposes that basal layer cells have equivalent long-term renewal potential, with stochastic division and differentiation decisions driving clonal expansion and loss through neutral competition.^8–14^ In this case, there is little deterministic signal, but rather a balanced set of decision probabilities downstream of a fluctuating signal. Although these models provide different explanations for the observed heterogeneity in clonal behavior, neither establishes the underlying signals that generate variability in individual cell fate decisions.

**Figure 1.**
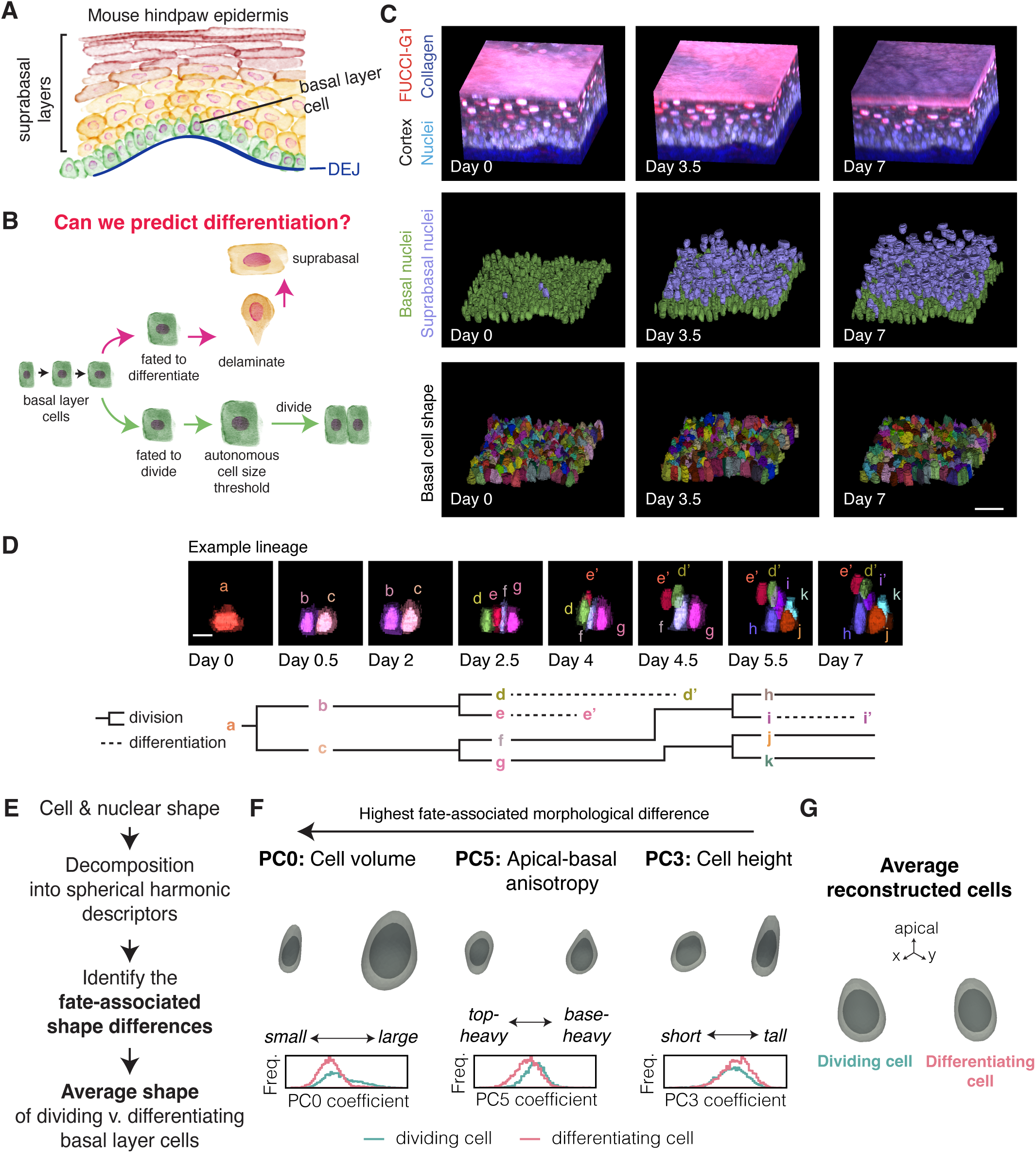
Dense lineage tracking reveals the morphological dynamics of epidermal basal layer cells during cell division and differentiation. A. The mammalian epidermis is composed of multiple layers of keratinocytes, with epidermal stem cells residing in the basal layer. B. Basal layer cells continually decide to divide or differentiate by delaminating into the suprabasal layers. C. Dense reconstruction of the fate, morphology, and descendants of all basal layer cells across 7 days using intravital imaging data from Mesa, Kawaguchi, & Cockburn, *et al*. (2018). Top: 3D volumetric rendering of hindpaw tissues expressing *H2B-Cerulean* (nuclei), *FUCCI2* (G1 cell cycle), *GFP-Actin* (cell cortex outline). Collagen imaging second harmonic generation (SHG) was used to identify the DEJ. Middle: Descendants of initial basal layer cells were tracked. Basal layer nuclei are rendered in green. Suprabasal layer nuclei are rendered in violet. Bottom: The shapes of all basal layer cells were segmented from the cell cortex. Each individual cell is rendered in a random color. Scale bar is 25 µm. D. Example basal layer lineage followed from day 0 to day 7. Each basal layer cell is rendered as a solid nucleus and translucent cell volume. Each suprabasal layer cell is shown as a solid nucleus only. The reconstructed lineage tree is shown. Scale bar is 10 µm. E. Strategy to identify the morphological differences between dividing and differentiating basal layer cells. F. The principal components (PC) of basal layer morphology were ranked by those that were most different between the cells that divided and the cells that differentiated. The top three ranked PCs are: PC0 (cell volume), PC5 (apical-basal anisotropy), and PC3 (cell height). The corresponding representative cell and nuclear morphologies are shown. The corresponding PC coefficients for dividing or differentiating cells are also shown. G. Reconstructed cell and nuclear morphologies of the average dividing and differentiating basal layer cell.

A major challenge in understanding cell fate heterogeneity is distinguishing inherently stochastic decisions from those biased by molecular or environmental factors. Traditional approaches based on population measurements, single-cell transcriptomics, or fixed-tissue imaging can reveal differences between cells but cannot establish whether these differences precede or follow their fate decisions. Longitudinal single-cell imaging offers a powerful alternative. By directly observing the eventual fate of individual cells, one can retrospectively examine their earlier molecular, morphological, and environmental characteristics to identify features that predict subsequent behavior. We and others have used this approach to identify precise molecular thresholds or metabolic states that govern cell cycle commitment in budding yeast and mammalian cells, demonstrating that seemingly variable cell behaviors can be highly predictable from their underlying molecular states.^15–17^

Applying this approach within adult tissues such as the skin is particularly challenging because individual cells must be tracked over extended periods while simultaneously reconstructing changes in their morphology and local environment. Nevertheless, longitudinal intravital imaging of the mammalian epidermis has begun to reveal how cell-intrinsic regulatory mechanisms interact with local tissue dynamics. Previous work showed that division and differentiation are locally coordinated, with the delamination of one basal layer cell promoting the subsequent division of a neighbor.^12^ Our previous studies demonstrated that heterogeneity in cell cycle duration can emerge from the interaction of cell-autonomous size control with local growth conditions.^18,19^ Basal layer cells must grow beyond a size threshold to enter S phase, while their growth rates depend on local cell context. Thus, differences in cell cycle duration can arise without requiring distinct stem cell subtypes. However, while these mechanisms help explain when cells divide, they do not explain why some cells continue to proliferate while others differentiate, or whether their subsequent fates can be predicted from features of their local tissue environment.

Here, we addressed the question of why some stem cells divide and others differentiate. To do this, we combined longitudinal *in vivo* imaging with 4D image analysis to reconstruct the morphological and fate histories of basal layer cells within the mouse hindpaw epidermis over one week. We tracked individual differentiating and dividing lineages and asked whether features of each cell’s local microenvironment could predict its subsequent fate or that of its descendants. Interpretable machine learning models identified local DEJ curvature as an unexpected predictor: valley regions favored continued proliferation and basal-layer retention, whereas ridge regions favored delamination and differentiation. Consistent with this spatial bias in short-term fate, valley and ridge regions were enriched for stem cell-associated Itga6 expression and the differentiation-associated *Krt10* expression, respectively. Moreover, clonal lineage tracing showed that cells initially in valleys proliferated more than cells in ridges. DEJ geometry remained remarkably constant over many weeks, substantially longer than the lifetime of an individual basal layer cell. Similar effects were seen in human epidermal equivalents grown on devitalized dermis, indicating that living dermal cells are not required for curvature-based spatial patterning. Together, our results support a model in which stable DEJ topography physically organizes the epidermal stem cell niche by providing spatial information that patterns individual fate decisions.

## Results

### Longitudinal imaging reconstructs epidermal cell fate and morphology across generations

To investigate the cellular and environmental dynamics underlying division and differentiation, we reconstructed the lineages and morphological trajectories of all basal layer cells initially present within two approximately 100 μm × 100 μm × 80 μm volumes of mouse epidermis. This gave complete lineage trees of 539 basal layer cells present at day 0 and whether they or their daughters divided or delaminated. We quantified the 3D nuclear and cellular shapes for every basal layer cell and the 3D nuclear shape of every suprabasal layer cell (**Figure 1C; Figure S1A-B**). We then integrated these temporal lineage trees with spatial maps of cell-cell contacts to generate a spatiotemporal tissue graph capturing the morphology, local environment, and fate history of each cell (**Figure 1D; Supplemental Figure S1C-E**).

To validate our lineage reconstruction, we compared our measured tissue-dynamic parameters with estimates from previous studies. The observed distribution of cell-cycle lengths (71.3 ± 23 h, mean ± SD) was consistent with estimates derived from label-dilution experiments (3 days).^13^ Moreover, only one basal layer cell did not either divide or differentiate during the seven-day imaging period, consistent with previous evidence that long-lived label-retaining cells may be rare in the basal layer (**Supplemental Figure S1E**).^13,20^ We next quantified daughter-cell fate outcomes from the reconstructed lineage trees. Among 303 divisions for which the fates of both daughters could be determined, 72 (24%) produced two dividing daughters, 93 (30%) produced two differentiating daughters, and 138 (46%) produced daughters with asymmetric fates. Our results are statistically consistent with the fraction of asymmetric outcomes reported in a previous study (*p* = 0.5; unpaired two-tailed t-test).^11^

To compare our morphodynamic measurements with known morphological features of epidermal stem cells, we parameterized the 3D cellular and nuclear morphology of each cell using spherical harmonics and performed principal component analysis on the resulting coefficients (**Figure 1E; Supplemental Figure S2A**).^21^ Reconstruction of the principal components (PCs) produced visually interpretable modes of shape variation (**Supplemental Figure S2B**). The modes that differed most strongly between cells that subsequently divided and those that differentiated mainly captured variations in cell volume (PC0), apical-basal anisotropy (PC5), and cell height (PC3) (**Figure 1F; Supplemental Figure S2C-F**). The average dividing cell was therefore larger, more pyramidal, and slightly shorter than the average differentiating cell (**Figure 1G**). These differences are consistent with our previous work showing that basal layer cells must cross a cell size threshold before entering S phase and previous observations that delaminating cells gradually lose basal contact area during their last cell cycle.^14,19^ Thus, our 4D morphodynamic dataset recapitulates the established features of basal layer division and differentiation while resolving their dynamics across individual lineages.

### Mother cell morphology and local environment predict daughter cell fate

To determine how early fate-associated differences emerge, we analyzed mother cells that produced daughters with known subsequent fates: both daughters divided, both daughters differentiated, or one divided and the other differentiated. Previous work has shown that basal layer cells expressing differentiation markers, including keratin 10 (Krt10), can still divide, indicating that differentiation-associated changes can precede cell cycle exit.^14^ We therefore asked whether mother cell morphology contained information about the subsequent fates of its daughters (**Figure 2A**). We also quantified features of each mother cell’s local environment to determine whether tissue context provides additional information about the subsequent fates of its daughters (**Figure 2B; Supplemental Table S1**).

**Figure 2.**
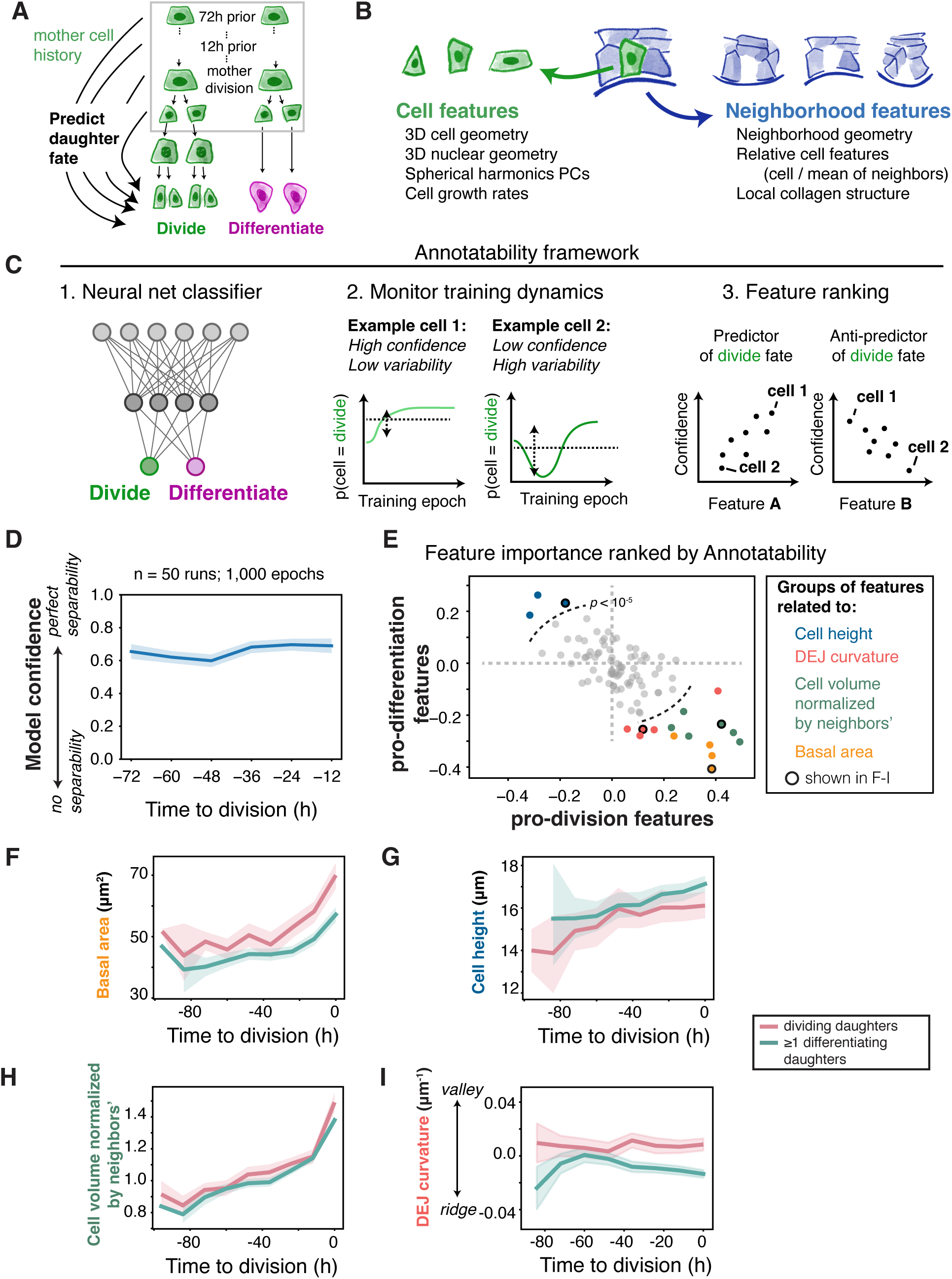
Interpretable machine learning models reveal the cell morphological features of the mother cell and microenvironment that predict daughter cell fate. A. Schematic of predictive modeling framework. After aligning mother cells by their division frame, features of the mother cell and its microenvironment were used to predict the eventual fate of daughter cells. Dividing daughters refer to daughter pairs that both went on to divide further. Differentiating daughters refer to daughter pairs where at least one daughter differentiated. Features measured prior to cell division were used to predict subsequent cell fate. B. Cell and microenvironment features extracted from imaging data were used for predictive modeling (see **Methods**). C. Annotatability framework was used to determine morphological features important for prediction accuracy. In this framework, a multilayered perceptron network was used to predict the daughter cell fate based on input mother cell features. The in-model probability (of either fate) of each cell was quantified over the training epochs. Model confidence is the probability mean, while model variability is the probability’s standard deviation. Features that correlate or anti-correlate highly with model confidence are features that contain information important for discriminating between the cell fates. D. The mean model confidence was quantified across the mother cell cycle. E. Spearman’s correlation between feature value and model confidence is averaged across all mother cell time points. Average feature correlation with confidence in division (pro-division) and differentiation (pro-differentiation) are plotted against each other. Four coherent groups of features were identified among those that correlate highly with model confidence in either division or differentiation: basal area, cell height, DEJ curvature, and relative cell size. Features highlighted in F-I are circled in black. Dotted line is a threshold of P = 1e-5 (see **Methods**) F-I. The dynamics of the indicated mother cell feature is shown for those that give rise to two dividing daughter cells or at least one differentiating daughter cell.

To predict whether daughter cells divide or differentiate, we trained a neural network classifier using the mother cell’s history. We grouped mothers producing daughters with asymmetric fates together with those producing two differentiating daughters. Hereafter, we refer to these two classes as division-biased and differentiation-biased mothers, respectively. We examined the model’s learning dynamics using Annotatability, a method that measures how confidently and consistently individual samples are learned over successive training epochs to quantify data separability and identify discriminating features (**Figure 2C**).^22^ For each cell, the mean of its classification probability represented the model’s confidence in its inferred label. Aggregate confidence across all cells therefore provides a measure of the separability of the two fate classes from the input features.

We applied this Annotatability framework to all dividing mothers for which the subsequent fates of both daughters were known. Mother-cell trajectories were aligned by their time of division, and separate models were trained using snapshots obtained at progressively earlier time points. After accounting for differences in sampling across time points, model confidence remained unexpectedly stable from immediately before division to 72 h before division, albeit at a moderate confidence of ∼0.65 (**Figure 2D**). For all time points, the model was more confident in its classification of division-biased mother cells compared to differentiation-biased mother cells (**Supplemental Figure S3A**). We also trained a random forest classifier on the same data and found that it also achieved consistent accuracy across mother cell history (**Supplemental Figure S4A-C**). Thus, features associated with future daughter-cell fate are already detectable in the mother cells and their microenvironments at least three days prior to division, indicating that a fate bias emerges before the final mother-cell cycle.

To determine the specific features that were most discriminating in classifying daughter fates, we used Annotatability to examine the learning dynamics of the neural net models. We correlated the mother cell features with model confidence, which quantifies how much discriminatory power they contain (**Supplemental Figure S3B-C**). These features formed coherent groups of pro-division and pro-differentiation predictors (**Supplemental Figure S3D-E**). The highest-ranking cell-intrinsic features included basal area and cell height, whereas the leading microenvironmental features included cell volume relative to neighboring cells and local DEJ curvature (**Figure 2E**). Differences in basal area and height resembled those observed in differentiating daughters: differentiation-biased mothers had smaller basal footprints and were taller than division-biased mothers, indicating that mother cells already acquire the morphological features associated with differentiation (**Figure 2F-G**). Our analysis also revealed fate-associated differences in the local cellular environment that were previously unknown. Although division- and differentiation-biased mothers had comparable absolute volumes (**Supplemental Figure S3F**), differentiation-biased mothers tended to be smaller than their immediate neighbors (**Figure 2H**). Unexpectedly, local DEJ curvature also emerged as a strong predictor of daughter-cell fate (**Figure 2I**). Consistent with the neural network classifier, the random forest classifier also identified basal shape and DEJ curvature as predictive features (**Supplemental Figure S4D**). Together, these findings show that differentiation-associated morphological features emerge during or prior to the mother cell division, and identify relative cell size and DEJ curvature as previously unknown features of the microenvironment associated with subsequent daughter fate.

### DEJ curvature correlates with epidermal stem cell shape and subsequent fate

The DEJ of the mouse interfollicular epidermis has generally been regarded as relatively flat, and its microscopic undulations have not been considered biologically significant.^23,24^ We therefore investigated the unexpected finding that local DEJ curvature predicted daughter cell fate. We defined positive curvature as concave, valley-like regions of the epidermis that extend into the underlying dermis, and defined negative curvature as convex, ridge-like regions that project away from the dermis (**Figure 3A**). We first asked whether these valley and ridge regions were associated with differences in the morphology of basal layer cells, which were more robustly associated with differentiation in our machine learning models. We found that DEJ curvature and cell morphologies all covaried, with cells residing in valley regions having larger basal areas and shorter cell heights than cells residing in ridge regions (**Figure 3B-C**).

**Figure 3.**
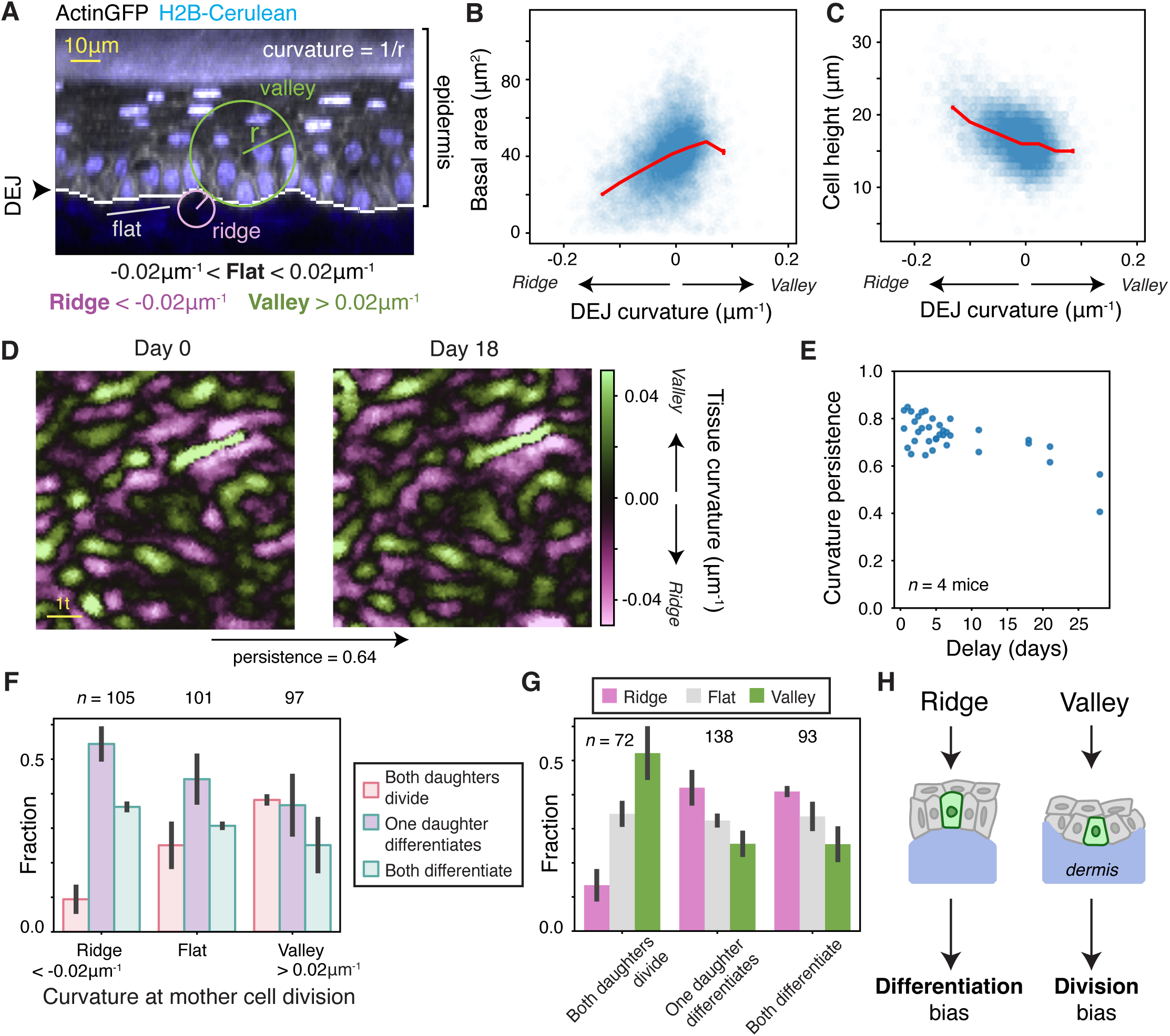
Basal layer cell geometry and fate are associated with long-term differences in DEJ curvature. A. Example illustration of DEJ curvature definition. An orthogonal slice of the epidermis expressing H2B-Cerulean and GFP-Actin is shown. The reconstructed DEJ is shown as a white line. Curvature is defined as the reciprocal of the radius of the sphere that best fits the local surface (see **Methods**). B-C. Cell geometry is correlated with DEJ curvature differences. Basal area (B) and cell height (C) plotted against the local DEJ curvature. D. DEJ curvature is stable over many weeks. *In vivo* imaging was used to estimate the DEJ curvature of the same tissue region on the mouse hindpaw at 18 days apart. After image registration, the maximum of the spatial correlation of the DEJ curvature between the two time points was used to define its persistence over time. E. The DEJ curvature persistence was quantified for 4 mice at different intervening times as indicated. F. The daughter cell fates for mother cells residing at ridge, flat, or valley regions of the DEJ at the times of their division. G. The DEJ curvature of mother cells giving rise to dividing, differentiating, or asymmetrically fated daughter pairs. H. Schematic showing that long-term ridge and valley regions of the DEJ preferentially lead to stereotyped cell geometries to pattern basal layer fate.

Since cell geometry and DEJ curvature covaried, we next asked whether differences in DEJ curvature are caused by the transient behaviors of individual basal layer cells, or are instead persistent structural features of the tissue. To determine this, we used longitudinal imaging to repeatedly image the same regions of the hindpaw epidermis at varying intervals and calculated the correlation between DEJ curvature maps across the two timepoints, which we termed persistence **(Figure 3D; Supplemental Figure S5A-C**). We found that the DEJ curvature was remarkably stable from day to day, showing appreciable remodeling only after approximately 20 days (**Figure 3E**). Thus, the local geometric landscape of the DEJ persists across multiple generations of basal layer cells, making it unlikely that its spatial pattern simply reflects the transient morphology or behavior of individual basal layer cells.

We next quantified the relationship between DEJ curvature and subsequent daughter cell fate. To do this, we binned the DEJ curvature underlying each mother cell into ridge (< -0.02 µm⁻¹), valley (>0.02 µm⁻¹), or flat regions and calculated the joint probability of a mother cell’s local curvature and number of differentiating daughters it produced (**Figure 3F-G; Supplemental Figure S5D**). The relationship between curvature and fate was asymmetric: mothers whose daughters both divided were preferentially found in valleys and depleted from ridges, whereas mothers producing at least one differentiating daughter showed a much weaker preference for ridges over valleys. These results identify DEJ curvature as a persistent feature on the tissue length scale that is associated with subsequent cell fate.

### DEJ curvature is associated with differentiation gene expression

Next, we asked whether the spatial association between DEJ curvature and cell fate is also reflected in the molecular state of basal layer cells (**Figure 4A**). To do this, we first quantified the promoter activity of *Krt10*, a well-established marker of epidermal differentiation (**Figure 4B**). We used a *K10-rtTA; pTRE-H2B-GFP* mouse line in which a transgenic *Krt10* promoter drives the expression of a nuclear marker following 3 days of doxycycline treatment. Previous work has shown that this reporter accurately reflects *Krt10* transcription.^14^ Using intravital imaging, we measured the spatial pattern of *Krt10* promoter activity while simultaneously reconstructing the DEJ and calculating its local curvature (**Figure 4C-D**). We then generated computationally flattened images of the basal layer to visualize the pattern of *Krt10* promoter activation alongside DEJ curvature (**Figure 4E**). Consistent with the fate bias observed in longitudinal imaging, we found that *Krt10* promoter activity correlates with DEJ curvature, with ridge regions having the highest reporter intensity (**Figure 4F; Supplemental Figure S6A-D**).

**Figure 4.**
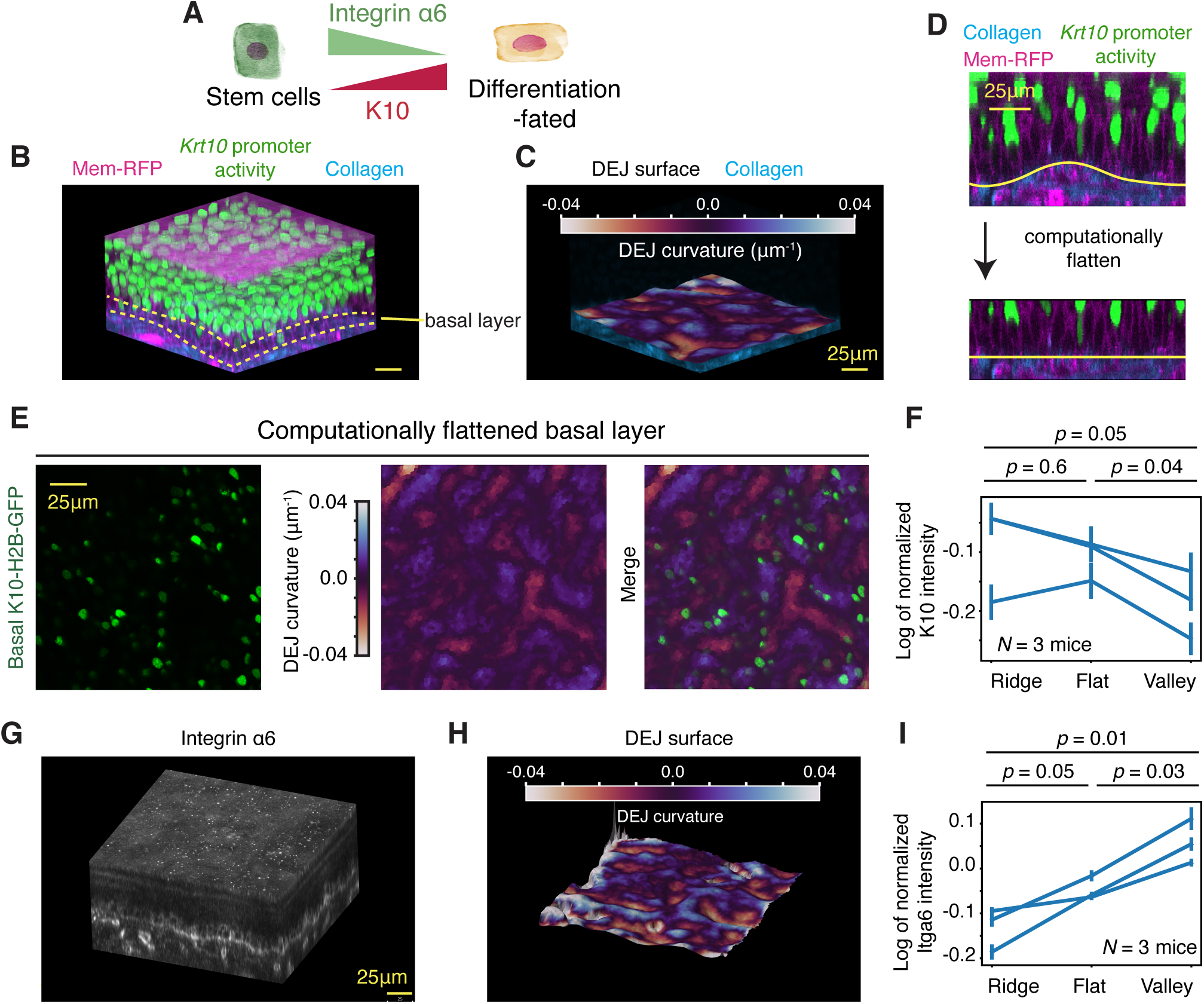
DEJ curvature is associated with molecular markers of basal layer stemness and differentiation. A. Integrin α6 was used as an epidermal stem cell marker, while *Krt10* promoter activity was used as a differentiation marker. B. Mouse hindpaw epidermis expressing *Krt10* promoter activity reporter and membrane-RFP was imaged in 3D. For visualization, the basal layer is identified using yellow dotted lines. C. The DEJ surface was reconstructed from the SHG collagen signal. The surface was rendered as a 3D mesh model. The surface color shows the local DEJ curvature. D. The 3D z-stack was computationally flattened using the reconstructed DEJ surface model and used to quantify basal *Krt10* promoter activity intensity. E. The computationally flattened basal *Krt10* promoter activity signal is shown alongside DEJ curvature. F. The log of mean-normalized *Krt10* promoter activity intensity is shown for ridge, flat, and valley regions. N = 3 mice. Data from individual mice are connected. G. Itga6 immunofluorescence signal from whole-mount full-thickness hindpaw skin is shown as a rendered 3D volume. H. The DEJ surface was reconstructed from the 3D Itga6 signal. The surface was rendered as a 3D mesh model. The surface color shows the local DEJ curvature. I. The log of mean-normalized Itga6 intensity is shown for ridge, flat, and valley regions. N = 3 mice. Data from individual mice are connected. The p-values are calculated using two-sided paired Student’s *t*-test.

We next asked whether there is a pattern complementary to the differentiation marker *Krt10* for a marker associated with stemness. To test this, we chose to examine protein expression of integrin α6 (Itga6), a basal layer marker that binds to components of the basement membrane along the DEJ (**Supplemental Figure S6E**). A previous single cell transcriptomic analysis of epidermal differentiation trajectories showed that *Itga6* expression decreases during differentiation.^14^ We used whole mount immunofluorescence to quantify the spatial pattern of Itga6 expression in intact, full-thickness skin tissues (**Figure 4G-H)**. We found that in contrast to *Krt10*, Itga6 expression was highest in the valley regions, consistent with our live-cell imaging results showing that cells in valleys were more likely to divide (**Figure 4I; Supplemental Figure S6F**). Thus, ridge and valley regions of the DEJ are associated not only with different subsequent lineage behaviors but also with opposing spatial patterns in stem- and differentiation-associated marker expression.

### DEJ curvature is associated with long-term clonal expansion

We have shown that DEJ curvature is associated with the fate decisions of individual cells and with the spatial distribution of differentiation markers. However, even modest differences in the probabilities of division and differentiation should accumulate over successive generations to produce substantial differences in clonal growth. Because DEJ geometry remains stable for weeks, we asked whether the initial local curvature of a basal layer cell is associated with the subsequent expansion of its lineage.

To test this, we first performed *in silico* lineage tracing using our reconstructed lineage graphs (**Figure 5A**). Over the seven-day imaging period, the initial DEJ curvature underlying each basal layer cell was associated with its subsequent clonal expansion, measured both by the number of basal descendants at day 7 and by the total number of descendants produced (**Figure 5B; Supplemental Figure S7A**). We next asked whether this relationship persisted over longer timescales. We used *K14CreERT; mTmG* mice to sparsely label basal layer clones with membrane-GFP and tracked their expansion relative to their initial local DEJ curvature (**Figure 5C**). Over 15 days, clones originating in valley regions expanded more than those originating in ridge regions (**Figure 5D**). Thus, the spatial biases in individual cell fate associated with DEJ curvature extend across multiple generations, producing persistent differences in clonal growth.

**Figure 5.**
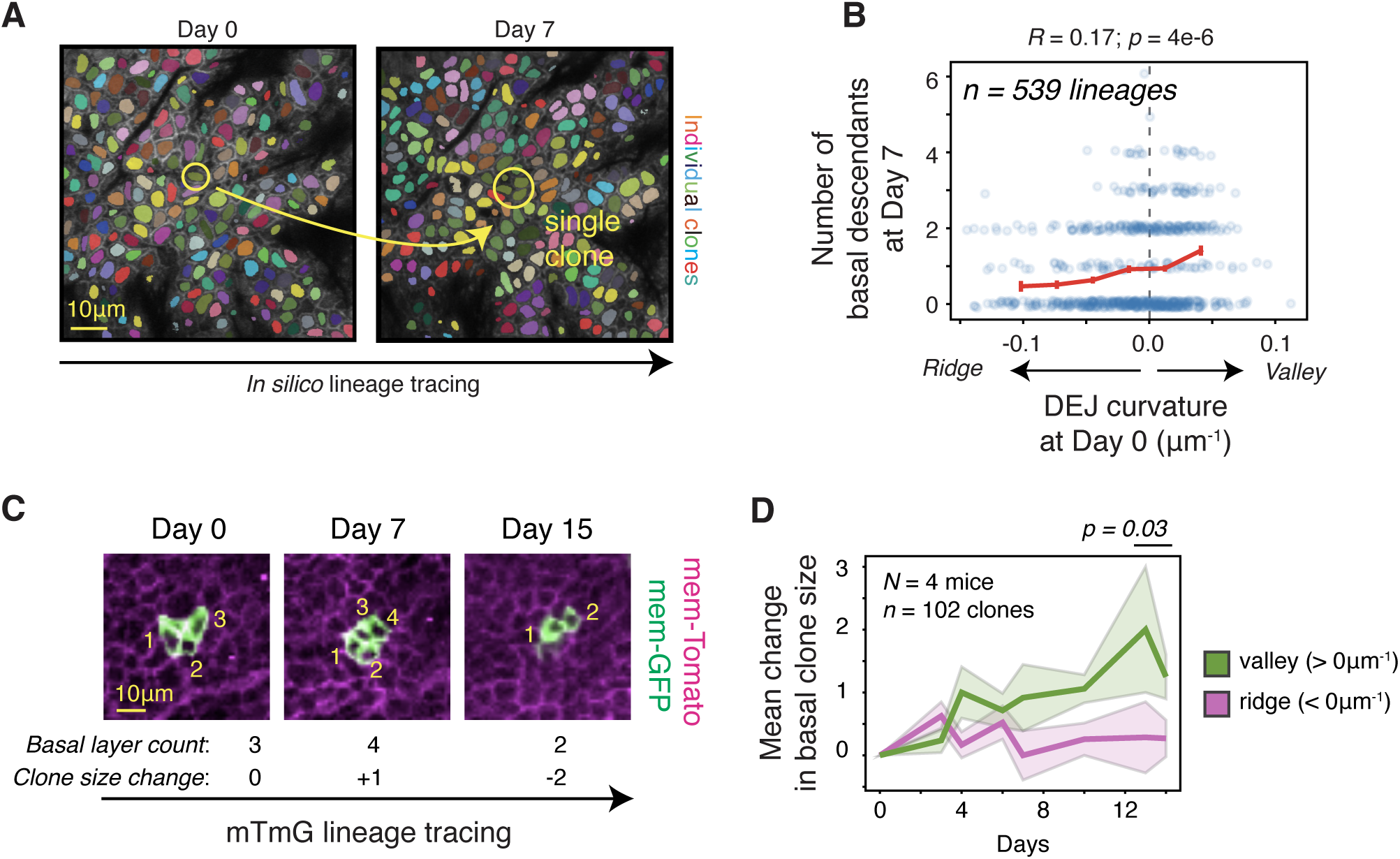
DEJ curvature is associated with differences in clonogenicity. A. Example of *in silico* lineage tracing using single cell tracking data. One olive-colored cell gives rise to 3 basal descendants. B. The number of basal descendants of a clone after 7 days is correlated with the DEJ curvature of its initial founding basal layer cell. R is Pearson’s correlation; the p-values were obtained against a null hypothesis of no correlation. C. Genetic lineage tracking using sparse *mTmG* clones. An example of a clone that was longitudinally quantified by counting changes in the number of membrane-GFP-positive basal layer cells within each clone. N = 4 mice. The p-values are calculated using two-sided unpaired Student’s *t*-test on clone-level statistics. D. The mean changes in basal clone size within ridge and valley regions are shown for the *mTmG* mice. Solid line is mean and shaded bars are standard error.

### Basal layer cells are preferentially retained in DEJ valleys in the absence of living dermal cells

The association between DEJ curvature and basal layer cell fate could arise through several mechanisms. Tissue geometry could directly influence basal layer cells through differences in cell packing and mechanical forces, or ridges and valleys could have different extracellular matrix (ECM) structures or compositions. Alternatively, DEJ curvature may covary with other features of the dermis that can locally signal to nearby keratinocytes, including proximity to lymphatic and blood vessels, fibroblasts, and immune cells. Distinguishing among these possibilities requires separating the relatively stable geometric and chemical features of the DEJ from dynamic signals arising from living cells in the dermis.

To determine whether living dermal cells are required for curvature-biased basal layer cell fate, we used human epidermal equivalent cultures.^25,26^ In this model, primary neonatal human keratinocytes are seeded onto devitalized human dermis and cultured at an air-liquid interface. Over six days, the keratinocytes form a stratified epidermis with appropriate layer-specific gene expression.^25^ We seeded 1-5% GFP+ keratinocytes as sparse tracer cells among GFP-keratinocytes. We hypothesized that the GFP+ tracers would initially occupy valleys and ridges with equal probability, but that preferential differentiation in ridges and continued division in valleys would progressively enrich GFP+ basal layer cells within valleys (**Figure 6A**). Two days after seeding, the average DEJ curvature underlying GFP+ tracer cells was close to zero and indistinguishable from random positions, indicating that GFP+ tracer cells were seeded uniformly across the dermis (**Figure 6B-C**). By day 6, after stratification, GFP+ basal layer cells were preferentially located in valley regions relative to random positions (**Figure 6C**). Therefore, curvature-biased retention of basal layer cells emerges in an organotypic human epidermis lacking living dermal cells, indicating that dynamic signaling from dermal cells is not required to establish this spatial bias.

**Figure 6.**
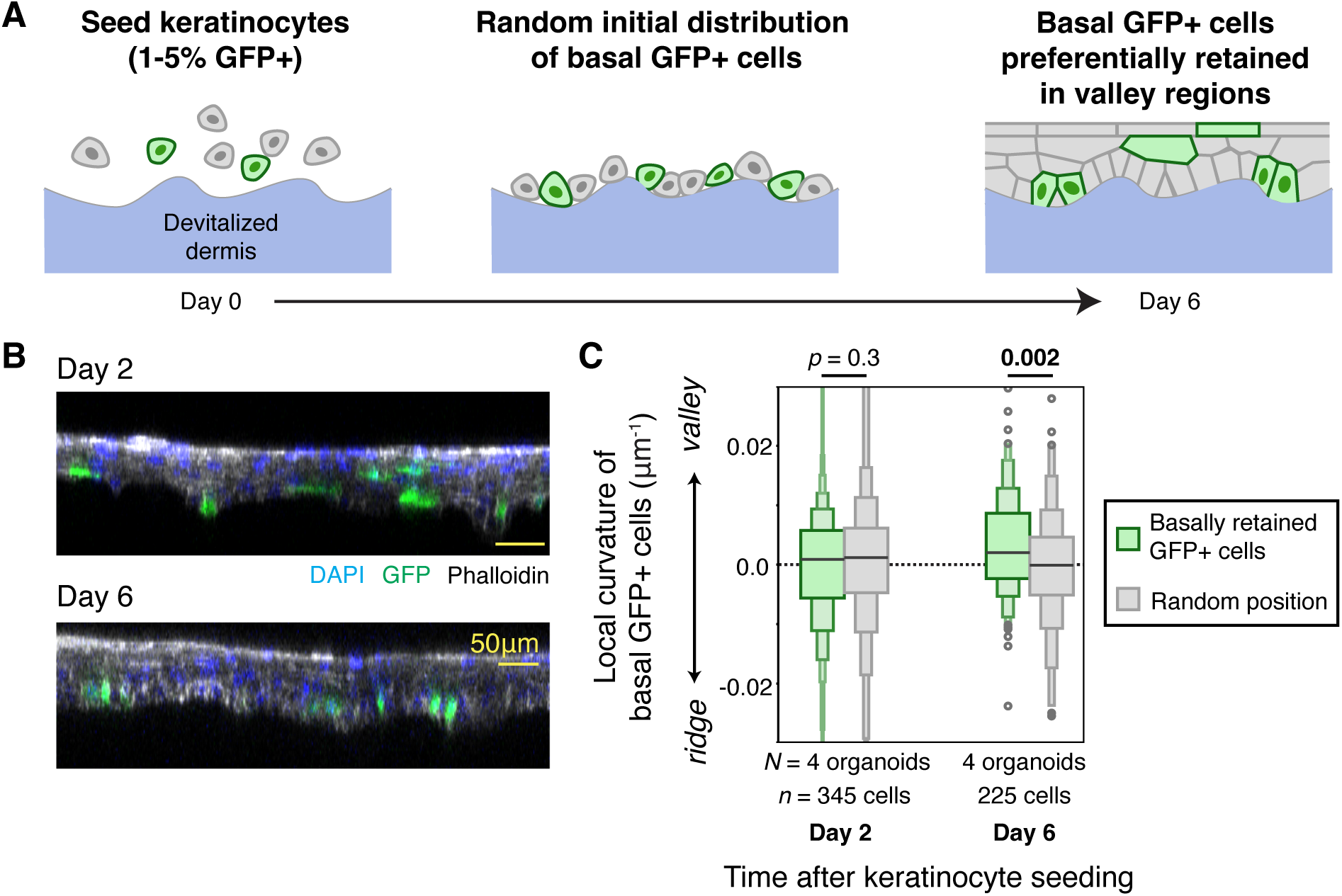
Living dermal cells are dispensable for DEJ geometry to bias basal layer cell fate. A. Human organotypic epidermis models were grown by seeding keratinocytes on top of devitalized dermis in an air-liquid interface device. A sparse population (1-5%) of GFP+ tracer keratinocyte cells were mixed with GFP-keratinocytes. The DEJ curvature underlying basal GFP+ cells was quantified during the maturation of the epidermal culture. B. Example orthogonal slices of epidermal organoids at 2 days and 6 days post keratinocyte seeding. C. The local DEJ curvature underlying basal GFP+ cells was quantified at 2 days and 6 days after keratinocyte seeding. For each timepoint, random spatial positions were also measured to quantify the full distribution of DEJ curvature within the field of view, which is densely occupied by the skin culture. The p-values are calculated using two-sided unpaired Student’s *t*-test on cell-level statistics.

## Discussion

Our results identify DEJ curvature as a stable feature of the epidermal stem cell niche that spatially patterns basal layer cell fate (**Figure 7**). Valley regions favor continued proliferation and basal layer retention, while ridge regions are associated with differentiation and delamination. Although individual basal layer cells turn over within days, local DEJ geometry persists for weeks and is associated with differences in cell morphology, differentiation marker expression, daughter cell fate, and long-term clonogenicity. These findings suggest that functional heterogeneity within the basal layer can emerge from larger scale differences in each cell’s multicellular environment. More broadly, this study reveals that features of the tissue’s physical landscape, however subtle, can still organize stem cell behavior.

**Figure 7.**
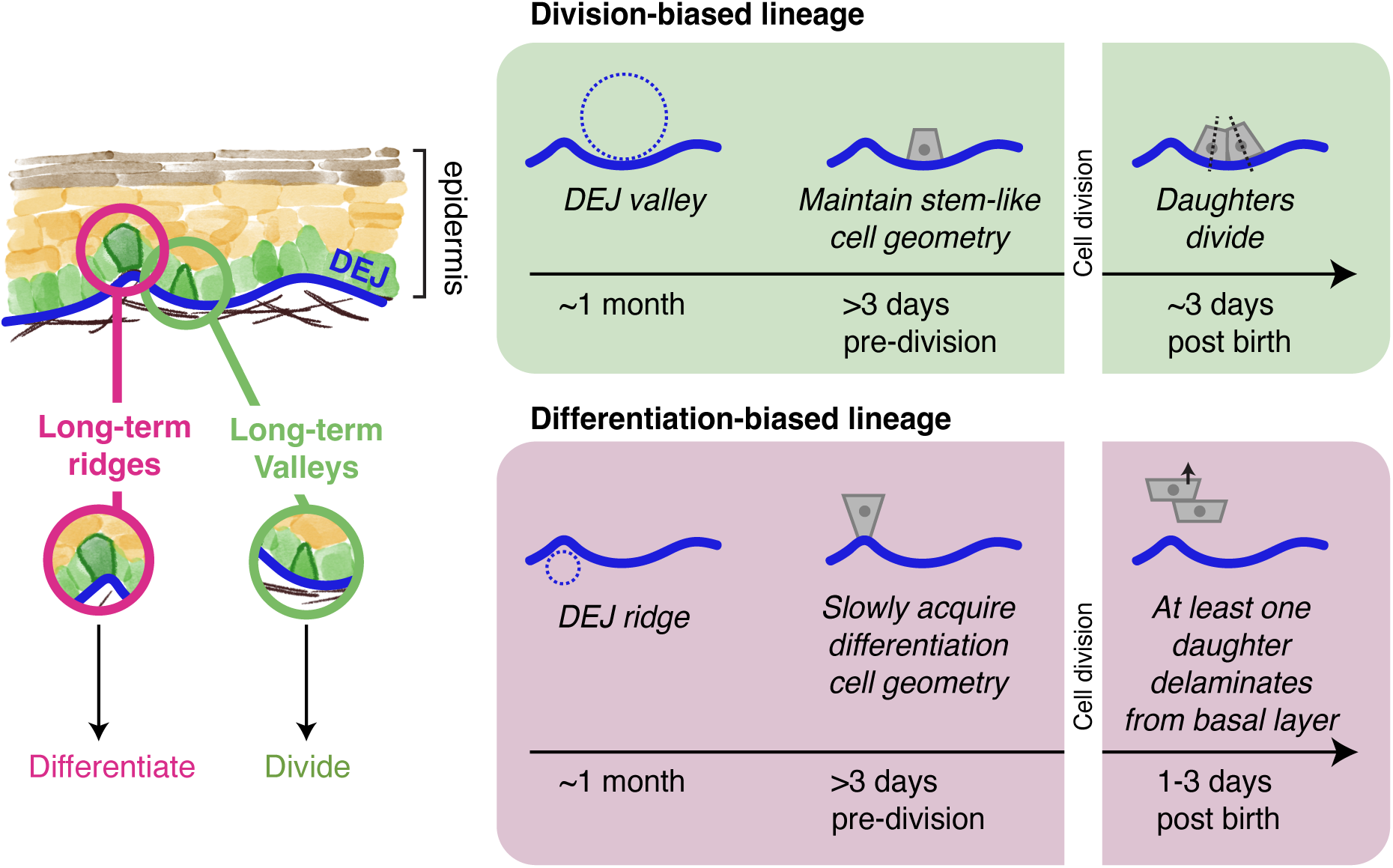
Schematic model indicating the effect of persistent tissue geometry on cell fate selection in the mouse epidermis.

Similar longitudinal imaging and computational analysis of cell dynamics have recently been used to identify features predictive of future cell fate, especially in cell culture models and developing and regenerating organs.^27–31^ In developing tissues, changes in individual cell behavior often coincide with broader changes in tissue organization, making it difficult to disentangle persistent predictors of cell fate from features associated with tissue remodeling. In contrast, the relatively stable architecture of the homeostatic adult epidermis allows us to identify a persistent feature of the tissue environment associated with cell fate.

Our findings provide a spatial dimension to ongoing models of stem cell heterogeneity in the murine interfollicular epidermis. Previous studies identified functional differences among basal layer cells, often interpreting them as evidence for distinct stem cell subtypes.^20,32–34^ Although our results do not distinguish between the equipotent and two-progenitor models, they suggest that differences in the microenvironment may be an important source of functional heterogeneity. However, we note that substantial variability in cell fate also remains unexplained by the morphological and environmental features we measured. Whether this residual variability reflects intrinsic stochasticity in molecular processes or additional, unmeasured determinants of cell fate, such as distinct progenitor identities or molecular states, remains an important open question.

Beyond the spatial organization of cell fate, our longitudinal observations also provide insight into the temporal dynamics of differentiation. We find that the subsequent fates of daughter cells can be predicted from the morphology and local environment of their mother cells up to three days before division. Thus, biases in cell fate emerge well before the daughters are born, suggesting that the propensity to differentiate develops gradually over time. Consistent with this view, our results and previous work show that basal layer cells can acquire differentiation-associated molecular and morphological features while retaining the capacity to divide.^14^ These observations suggest a model in which differentiation propensity changes gradually, but the eventual decision to leave the basal layer through delamination is discrete. Before this discrete transition, basal layer cells remain capable of proliferation, with entry into S phase dependent on reaching the cell size threshold identified in our previous work.^18,19^ Thus, cells can progress toward differentiation while continuing to grow and divide, suggesting that cell cycle progression and differentiation propensity are regulated by distinct but interacting processes. We propose that tissue architecture biases differentiation propensity, while the subsequent delamination of differentiating cells creates space for neighboring cells to grow and divide. In this view, epidermal homeostasis can emerge from feedback among tissue architecture, local coordination of cell behaviors, and cell-intrinsic size homeostasis.

The ability of subtle DEJ undulations to predict epidermal stem cell fate suggests a broader role for tissue geometry in organizing stem cell behavior across epithelia. In the small intestine, stem cells reside at the base of invaginated crypts, whereas most of their differentiating descendants move toward the villi, creating a pronounced spatial separation between regions of self-renewal and differentiation.^35^ Recent studies using intestinal organoids grown in engineered geometries have demonstrated that tissue curvature itself can contribute to this organization, with crypts emerging in regions of highest curvature.^36,37^ Human skin also exhibits prominent undulations formed by rete ridges and dermal papillae, which have long been implicated in the spatial organization of epidermal proliferation and differentiation, although the spatial locations and identities of epidermal stem cells within these structures remain debated.^24,38–40^ Whether these larger structures in human skin and the microscopic DEJ undulations observed in mice share similar organizing principles remains unknown.^41,42^ Our study and recent work on the undulating architecture of the mouse oral mucosa may begin to shed light on these connections.^43^ We propose that tissue geometry organizes stem cell behavior across a continuum of spatial scales and degrees of compartmentalization, from the pronounced separation of self-renewal and differentiation in the intestinal epithelium to the more subtle, probabilistic fate biases observed in the flatter mouse epidermis.

The mechanism by which DEJ curvature biases basal layer cell fate remains unresolved. Our human organotypic culture experiments suggest that signaling from living dermal cells is not required for this spatial patterning. One possibility is that fate biases arise from the distinct cellular morphologies adopted in valley and ridge regions. These curvature-associated cell morphologies could reflect altered basal tension, which mechanical models of the epidermis suggest is sufficient to bias cell fate.^44^ Morphological differences could also influence cell signaling by altering the areas of contact with neighboring basal layer cells, the underlying basement membrane, or overlying suprabasal layer cells. A precedent for such a mechanism comes from intestinal crypts, where the coupling between cell morphology and tissue curvature may enhance contact-dependent niche signaling between intestinal stem cells and neighboring Paneth cells.^45^ Our current experiments, however, cannot rule out differences in ECM or basement membrane structure and composition between valley and ridge regions that may separately influence basal layer cell fate.^46,47^ Direct manipulation of DEJ geometry independently of ECM composition will be necessary to determine whether curvature itself is an instructive physical cue for basal layer cell fate.

Our findings raise the possibility that changes in epidermal architecture may contribute to altered stem cell behavior in human disease contexts. If the relationship between DEJ geometry and cell fate observed in mice extends to human skin, changes in rete ridge architecture could alter the spatial distribution of proliferation and differentiation. This possibility is particularly relevant to aging, disease, and wound healing, where DEJ architecture undergoes substantial remodeling. During skin aging, rete ridges progressively flatten, accompanied by reduced epidermal proliferation.^47–50^ Conversely, psoriasis is characterized by pronounced elongation of rete ridges, excessive keratinocyte proliferation, and abnormal differentiation.^51–54^ Normal rete ridge architecture is also frequently lost following wound healing and scarring.^55^ Although these structural changes may arise as consequences of altered tissue dynamics, they could also actively contribute to dysregulated keratinocyte behavior. Indeed, restoring appropriate tissue topography has shown promise in maintaining stem cell function in engineered epidermis models.^56–58^ More broadly, our study suggests that changes in the physical architecture of stem cell niches may contribute to altered tissue homeostasis during aging and disease.

## Methods

### Mouse lines

All mice were handled in accordance with the guidelines of the Institutional Animal Care and Use Committee at Stanford University or Yale University. *K14-CreER* and *mTmG* mice were obtained from JAX. The *Membrane-GFP* mice were obtained from Dr. K. Hadjantonakis (Memorial Sloan Kettering Cancer Center). *K10-rtTA* mice were obtained from T. Lechler (Duke University). *pTRE-H2BGFP* mice were obtained from E. Fuchs (Rockefeller University). *K10-rtTA*; *pTRE-H2BGFP*; *mTmG* mice were given doxycycline (2 mg ml^−1^) in drinking water with 2% sucrose continuously, starting 3 days before imaging. All mice imaged in the study were 2-4 months old and never used for breeding. Mice of either sex were distributed randomly for experiments.

The mouse lines are listed in **Table 1**. The crosses are listed in **Table 2**.

**Table 1.** The mouse lines and crosses used in this study.

| **Mouse line** | **Catalog number** | **Source** | **Paper** |
|---|---|---|---|
| *K14CreERT2* (*K14-CreER*) | #038390<br>RRID:IMSR_JAX:038390 | JAX | <sup>59</sup> |
| *mTmG* | #007676<br>RRID:IMSR_JAX:007676 | JAX | <sup>60</sup> |
| *Membrane-GFP (CAG-GFP)* | #011106<br>RRID:IMSR_JAX:011106 | K. Hadjantonakis | <sup>61</sup> |
| *K10-rtTA* | N/A | T. Lechler | <sup>62</sup> |
| *pTRE-H2B-GFP* | #005104<br>RRID:IMSR_JAX:005104 | E. Fuchs | <sup>63</sup> |
| *K14-H2B-mCherry* | N/A | V. Greco | <sup>64</sup> |
| *K14-H2B-Cerulean* | N/A | V. Greco | <sup>12</sup> |
| *K14-GFP-Actin* | N/A | E. Fuchs | <sup>65</sup> |
| *pR26-FUCCI2* | MGI:5491593 | V. Greco | <sup>66</sup> |

**Table 2.** Mouse crosses used in this study.

| Mouse cross | Experiment |
|---|---|
| *H2B-Cerulean; K14-GFP-Actin; FUCCI2* | Dense 4D reconstruction |
| *K14CreERT; mTmG* | Lineage tracing<br>DEJ persistence |
| *Membrane-GFP; FUCCI2* | DEJ persistence |
| *K10-rtTA; pTRE-H2B-GFP; mTmG* | *Krt10* promoter activity |
| *K10-rtTA; pTRE-H2B-GFP; mTmG; K14-H2B-mCherry* | *Krt10* promoter activity |

### Longitudinal intravital imaging

Dense annotations of basal layer cell lineage dynamics were based on a previously acquired longitudinal intravital imaging time course of *H2B-Cerulean; K14-GFP-Actin; FUCCI2* mice, where hindpaw images from an eight-week-old mouse were acquired at 12h intervals for 7 days. Two fields of view were used, consisting of 539 tracked lineages. Imaging of DEJ persistence, *Krt10* promoter activity, and *mTmG* genetic lineage tracing for this study were performed based on protocols previously described.^19^ During imaging, the mouse was kept under anesthesia by isoflurane inhalation through a nose cone and kept warm by a rodent heating pad. The mouse paw skin was immobilized between a custom-made stage arm and another stage arm holding a coverslip. The coverslip pressure was kept to a minimum to immobilize the tissue without deforming it. Images were taken on a Prairie View Ultima IV (Bruker) scope equipped with a DeepSee II tunable laser (Coherent) or a LaVision TriM Scope II (LaVision Biotec) laser scanning microscope equipped with both a Chameleon Vision II and Discovery two-photon lasers (Coherent). The sample was imaged with either an Olympus XLUMPLFLN 20x water immersion lens (NA = 0.95) or a Nikon water immersion lens (NA = 1.15). Images were acquired at 0.5-1 µm steps to image a total depth of ∼80 µm of tissue, covering the entire thickness of the epidermis. The GFP and tdTomato in *mTmG* mice were visualized simultaneously, using the excitation laser tuned to 920 nm and both a 525/50nm and a 595/50nm bandpass filter. For imaging *Krt10* promoter activity sensors alongside *mem-tdTomato*, the GFP was imaged using the Vision laser tuned to 920nm while tdTomato was imaged using the Discovery laser tuned to 1080nm. Collagen fibrils were visualized with a 460/50nm bandpass filter at 920nm or 940nm excitation.

To find the same tissue regions over time, cartographic notes were taken noting the idiosyncratic features of the skin itself, including folds and blood vessel patterns. The exact matching tissue region was confirmed by image registration using the underlying dermal collagen signal from session to session.

### Segmenting and tracking single cells in longitudinal skin images

A custom image analysis pipeline was built to register, segment, and track epidermal stem cells in the basal layer, based on previous work.^19^ In brief, the longitudinally acquired images were first registered using the second-harmonic generation (SHG) collagen signal, since collagen fibrils do not move during skin homeostasis. Cross-correlation was used to identify the respective z-positions between two time points that corresponded to each other, and a Python wrapper of StackReg was used to register the two tissue volumes.^67,68^ Cellpose 3.1.1.3 was then used to segment cells. 2.5D stitching mode was used, using the pretrained nuc and cell models and flow3D_smooth=0.^69^

Mastodon/FIJI was used to track the migration and division of every basal layer cell that was initially visible at Day 0 of the time course.^70,71^ The tracking was seeded with initial semiautomated tracking and manually corrected to reconstruct the lineage connection between all mother-daughter and sister cells. Each cell was labeled as basal or suprabasal based on whether it had a visible basal footprint (**Supplemental Figure S1A-B**). The coordinates of the lineage tracks were mapped onto the registered movie, and the corresponding nuclear and cell segmentations were collated. A custom napari workflow was used for manually inspecting, curating, and editing the 3D segmentations.^72^ Cell cycle timings were manually annotated based on the *FUCCI-Cdt1-mCherry* cell cycle reporter signal. A final script was used to collate the segmented and tracked cells to generate time-series measurements for every tracked epidermal basal layer cell lineage (**Supplemental Figure S1C-E**).

### Principal component analysis of cell morphology variation

3D cell and nuclear shapes were parametrized using spherical harmonics decomposition, using aics-shparam.^21^ Each cell or nuclear mask was first aligned to a shared XY planar orientation, since the XY laboratory coordinate is arbitrary. However, cells were not aligned by their axial orientation since the z-axis is aligned with the apical-basal axis of the tissue. Cells were fitted to spherical harmonics basis functions, up to Lmax=5. Then, all nuclear and cytoplasmic spherical harmonics coefficients were collated for PCA decomposition, which was implemented by scikit-learn.^73^ We focused our analysis on the first 9 PCs, which together explain >75% of all shape variation (**Supplemental Figure S2A**).

To reconstruct visual representations of the spherical harmonics PCs, we back transformed the loading weights of each PC into the original nuclear and cell spherical harmonics coefficients, as calculated by:

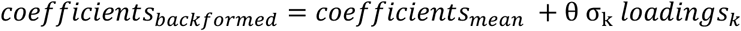

where *coefficients_mean_* is the mean harmonic coefficients across all samples, σ_k_ the square root of the variance explained by the k^th^ PC, *loadings_k_*the decomposition weights of the k^th^ PC onto harmonics coefficients, and θ the parametric variable representing the coordinate along the k^th^ PC. Using aics-shparam and these backformed coefficients, we reconstructed 3D mesh models corresponding to the 5^th^ and 95^th^ percentile values observed along that PC. This resulted in visually interpretable shape modes ranked by their relative contribution to overall variation (**Supplemental Figure S2B**).

### DEJ reconstruction and tissue curvature quantification

Two different methods were used to detect and reconstruct the DEJ, depending on the structure of the DEJ signal. The first method detects the sudden appearance or disappearance of image signal crossing the DEJ, for example the disappearance of epidermal signals from reporters expressed from a K14 promoter, or the appearance of SHG signal from dermal collagen. The second method detects signal like Itga6 staining, which are enriched at the DEJ.

The first method was based on intensity gradient changes using methods similar to prior work.^12,19^ Keratinocyte or collagen signals were blurred with a Gaussian filter with kernel size corresponding to ∼10µm in each direction. To detect the tissue boundary, the first derivative is taken along the Z-axis for each XY point. The boundary is defined as the z-slice at which the first derivative is maximized. The sign of the derivative is adjusted depending on the direction of imaging and the nature of the signal.

The second method was used to detect the DEJ from Itga6 stained image stacks. The Itga6 signal has local maxima in z at the DEJ, but could have noisy signal elsewhere, especially in the dermal blood vessels. To identify a continuous surface, all local maxima in z were found for each XY position, and neighboring XY pixels were connected if their maxima were within a small window (2 μm) of each other. All connected graphs were found and sorted by total size. The best graph was manually selected and used to reconstruct the final DEJ.

Since both methods sometimes yield discontinuities across XY regions, these discontinuities are detected using the connected components algorithm. The largest contiguous surface is then patched using bilinear interpolation. The final surface is saved as a pixel image for visual inspection alongside the original tissue images. For all subsequent analysis, the reconstructed DEJ was visually inspected alongside the raw image, and any artifacts were masked out manually for quantifications.

To reconstruct a 3D model of the DEJ, we resampled the pixel heightmap by a factor corresponding to ∼2-3µm resolution and generated a mesh surface using the trimesh library.^74^ Mean and Gaussian curvatures were estimated using the discrete curvature estimation method based on ball-mesh intersections implemented in trimesh.^75^ Both measures were normalized by πr², the surface area within the neighborhood if it were locally planar. Mean and Gaussian curvatures were expressed in μm⁻¹ and μm⁻², respectively. Unless otherwise noted, the neighborhood radius was 5 μm, and DEJ curvature refers to mean curvature.

### Measurement of the cellular microenvironment

An image analysis framework based on our previous work was used to quantify cellular and microenvironmental features.^19^ Briefly, the geometric features of individual cells were calculated using both its nuclear and cell segmentations. For each basal layer cell’s nuclear and cell segmentation, regionprops from scikit-image was used to generate individual object statistics, including cell and nuclear shapes and positions.^76^ To analyze cell apical and basal area, cell segmentation masks were computationally flattened using the z-coordinate of the DEJ to re-index the image stack (**Figure 4E**). Apical and basal areas were the union of projected area of the 4 topmost and 4 bottommost slices, respectively. The DEJ surface was then reconstructed and used to measure local tissue geometry and ECM characteristics, as described above.

Characteristics of the collagen channel were inferred from a maximum intensity projection of the SHG signal within 5 μm below the DEJ. We calculated the eigenvalues of the structure tensor (outer product of the two gradients of the SHG image). The orientation of the fibrils was defined as the principal eigenvector, while the coherence of the collagen fibrils was defined as the normalized difference between the eigenvalues. Non-keratinocyte epidermal cells were manually annotated by their *FUCCI2-Cdt1-mCherry* nuclear expression and lack of *K14-GFP-Actin* expression.

To measure cellular statistics in relation to those of its neighbors, we also reconstructed the cell-cell contact matrix for each movie frame independently. The actin cortex channel was carefully examined to determine which cells shared a cell-cell interface and the information encoded in a 2D contact image. Then, each cell’s individual statistic was used to generate a ‘relative’ statistic by dividing its value by the mean value of all its neighbors. For example, a cell’s relative cell volume would be defined as its cell volume normalized by the average volume of all its next neighbors.

The features used in the model are presented in **Supplemental Table S1**.

### Data preparation for machine learning analysis

To prepare a dataset where mother cell features could be used to predict daughter cell fate, we first found all mother cells for which daughter cell fate could be empirically measured within the time course. Daughters were denoted to have (1) both divided, producing 4 granddaughter cells, (2) both delaminated, producing 2 suprabasal layer cells; or (3) one divided and one delaminated. Fate (2) and fate (3) were pooled for our exploratory machine learning models. The daughter cells were aligned by their birth frame, and mother cells were aligned by their division frame. Feature vectors corresponding to daughter birth, mother division, and 12 h increments prior to mother division, up to 72h prior, were collated. Due to the need to ascertain daughter fate, mother timepoints included in the analysis tended to be earlier within the observation window of the time course.

### Fate-learnability analysis using Annotatability

To assess how readily division fate can be learned from cell features, we applied Annotatability, which scores each cell by the training dynamics of a neural network trained to predict its label.^22,77^ Divisions were labeled as self-renewing (both daughters divided) or differentiating (at least one daughter differentiated), and each timepoint before division, comprising 12h up to 72h prior to division at 12h intervals, was analyzed separately, as described in the previous section. Missing values were set to zero and features were *z*-scored. A fully connected network (two hidden layers of 32 and 16 units, ReLU) was trained for 1,000 epochs with stochastic gradient descent (learning rate 0.001, momentum 0.9), using class-balanced sampling and loss weighting. For each cell, confidence and variability were defined as the mean and standard deviation, across epochs, of the predicted probability of its true fate.

### Subsampling and feature correlations using Annotatability

The analysis was repeated on 50 random subsets of 400 divisions per timepoint, and mean confidence and variability were summarized overall and separately for each fate. To identify features associated with fate learnability, we computed Spearman correlations between each feature and confidence in fate prediction for each cell. Significant correlations (discovery threshold of *P* < 1e-5) were ranked, and features were prioritized by the number of timepoints at which they appeared among the ten strongest positive or negative correlations. Associations were assessed using Spearman’s rank correlation, implemented with scipy.stats.spearmanr. Two-sided p-values were calculated using the asymptotic Student’s t approximation with n − 2 degrees of freedom, where n denotes the number of paired observations.

### Tree-based classification of mother cells

Statistical models were built using the Python libraries scikit-learn.^73^ For random forest classification, RandomForestClassifier from scikit-learn was used. To correct for class imbalance, the over-represented class (differentiation-biased mothers) were subsampled to the same sample size as the under-represented class. Training/test splits were constructed by 80-20% split. For each set of mother cell features, 1000 repeated model fits were trained, including class rebalancing and test/train splitting, and an average confusion matrix was generated (**Supplemental Figure S4A-B**). The ROC-AUC scores were also calculated for each mother cell time point (**Supplemental Figure S4C**). The impurity importance was quantified from scikit-learn’s in-model implementation and averaged across all independent models (**Supplemental Figure S4D**).

### DEJ curvature persistence measurements

To measure how the spatial locations of valleys and ridges in the DEJ can change, we longitudinally measured the dermal collagen of a small region on the mouse hindpaw over time. Both *Membrane-GFP* and *mTmG* reporter mice were used. We then calculated the mean curvature of the DEJ across different time points. We used StackReg to register each pair of DEJ curvature maps, and calculated the normalized spatial correlation between them. The persistence was reported as the maximum spatial correlation.

### Immunofluorescence staining of mouse hindpaw keratinocytes

Immunofluorescence staining was performed on whole mount full-thickness skin. Hindpaws were harvested from animals and the skin was dissected from the rest of the paw. The nonhairy plantar skin was cut out, flattened onto filter papers, and fixed for 2 hours in 4% paraformaldehyde. The tissues were then washed 5 times in PBS and incubated in permeabilization buffer (1%BSA, 2% Triton-X, 5% normal goat serum, 0.2% Fish gelatin) for 3 days shaking at 4C. To ensure complete antibody penetration, the tissues were incubated with primary antibody (1:200, eBioGoH3 (GoH3), Rat monoclonal, eBioscience #14-0495-82) for 7 days. The tissues were washed 5 times in PBS and incubated in secondary antibody (1:1000, Goat anti-Rat IgG (H+L) Cross-Adsorbed Secondary Antibody, Alexa Fluor™ 488, #A-11006) for 7 more days, alongside Hoechst 33342. The tissues were washed 5 times in PBS and mounted epidermis side down using Vectashield Plus Antifade Mounting Medium (Vector Labs #H-1900-2) and stored at 4C.

### Confocal microscopy of fixed tissues

Fixed whole mount skin samples or organotypic cultures were imaged in 3D using confocal microscopy on a Zeiss LSM980 with a Zeiss LCI Plan-Neofluar objective (25x/NA=0.8) or Zeiss LD LCI Plan-Apochromat (40x/NA=1.2).

### Image flattening and quantification of *Krt10* promoter activity and Itga6 protein expression

To quantify basal layer *Krt10* promoter activity or Itga6 protein expression alongside DEJ curvature measurements, stacks of confocal or two-photon images were computationally flattened. To quantify *Krt10* promoter activity, z-slices of the hindpaw skin from *K10-rtTA; pTRE-H2B-GFP; mTmG* or *K10-rtTA; pTRE-H2B-GFP; mTmG; K14-H2B-mCherry* were taken. A basal slice of the flattened membrane-tdTomato channel was used to segment individual basal layer cells (**Supplemental Figure S6A-C**). A maximum intensity projection spanning 5 μm above the DEJ was made of the *Krt10* promoter activity channel and used to quantify the *Krt10* activity. For each cell, the mean *Krt10* promoter activity intensity and the average DEJ curvature were calculated and correlated.

To quantify Itga6 protein expression, the DEJ was reconstructed from whole mount full thickness skin images of wild-type mice stained with immunofluorescence against Itga6. A maximum intensity projection of the Itga6 channel was calculated from 2 μm above and below the DEJ mesh model. This intensity value was normalized by the mean intensity within the whole field of view, and a log10 of the fold change calculated. The DEJ curvature was binned, and the log10-mean-normalized intensity values quantified for each bin.

### *In silico* and genetic lineage tracing

*In silico* lineage tracing was done in the densely reconstructed longitudinal imaging dataset. We quantified the local DEJ curvature of an initially basal layer cell against the total number of descendants or the total number of basal descendants. To check that this curvature-associated difference in clone dynamics is not inflated by sister cells having the same fate, we binned each imaging field of view into 20 μm x 20 μm regions, corresponding to approximately 3-4 cells in length. From each region, only one basal layer cell was randomly selected to be quantified in the spatially blocked statistic. Confirming that this clonal statistic is not overly influenced by sister-pairs, we observed the same correlation between DEJ curvature and basal descendant count in spatially separated clones (*R* = 0.18; *p* = 0.046; Pearson’s correlation and bootstrap test; **Supplemental Figure S7B-C**).

Hindpaw skin from *K14CreER*; *mTmG* mice aged 2-4 months old were used to track clone dynamics over time. Membrane-GFP clones were sparsely labeled by spontaneous membrane-GFP induction in the absence of tamoxifen. Clone-size changes were measured relative to the first observation, rather than to the time of reporter recombination, which was unknown. The same clones were imaged longitudinally every 3 days for approximately 2 weeks. Baseline labeling prevalence ranged from 0.2% to 0.5% labelled clones per 100 basal layer cells (**Supplemental Figure S7D**). We assumed that new recombination events were relatively infrequent during the observation period and interpreted changes in basal membrane-GFP cell counts within tracked clones as reflecting basal-cell dynamics. However, ongoing recombination could contribute to these changes and could itself depend on DEJ curvature.

The time course images were registered across time points and the changes in basal membrane-GFP cell count per clone were quantified over time manually. The initial DEJ curvature was quantified by averaging curvature value within 5 μm of the clone centroid at day 0. To quantify the total density of membrane-GFP clones, the total number of cells within a field of view was estimated by dividing the image size by the average basal area of a basal layer cell. Then, the total number of cells at day 0 was calculated for each image region and a clone prevalence obtained.

### Organotypic epidermal cultures

Primary human keratinocytes were isolated from fresh surgically discarded skin and cultured in a 1:1 mix of Keratinocyte-SFM (Life Technologies #17005-142) and Medium 154 (Life Technologies #M-154-500). Organotypic regeneration of human epidermis was performed as previously described.^78^ Briefly, 500,000 cells were seeded onto devitalized human dermis (NDRI) in keratinocyte growth media (KGM) at the air-liquid interface and allowed to grow for 2 or 6 days. GFP-expressing keratinocytes were generated by inducing with lentivirus from pLEX-Puro-EGFP, and GFP+ cells were mixed at a 1 or 5% ratio with GFP-cells prior to keratinocyte seeding.

### Whole mount analysis of organotypic epidermal cultures

At the indicated times post keratinocyte seeding, organotypic epidermal cultures were fixed for 2 hours in 4% paraformaldehyde or for 15 min in 10% neutral buffered formalin. They were then incubated with Hoechst (1:1000) and phalloidin-iFluor-647 (1:1000, AAT Bioquest #23127) in a permeabilization buffer (1%BSA, 2% Triton-X, 5% normal goat serum, 0.5% Fish gelatin) at 4C overnight. The tissues were washed 3 times and stored in PBS at 4C and were mounted epidermis side down on a Matek imaging dish during microscopy analysis. Confocal microscopy was used to tile large fields of view of the epidermal culture at 3 μm z-slices. A surface model of the DEJ was reconstructed from the phalloidin channel. Any artifacts of failed surface reconstruction were manually masked out. The center of each GFP+ cell was annotated manually in napari, and the XY location of each GFP+ basal layer cell was used to calculate the local DEJ curvature by averaging a 5 μm square area around the cell centroid. Random positions were obtained by sampling the same number of cells from the field of view at random. Positions landing in masked regions was not counted.

### Statistics and software

Pearson correlations and p-values were calculated using scipy.stats.pearsonr. Two-sided unpaired and paired Student’s t-tests were performed using scipy.stats.ttest_ind or scipy.stats.ttest_rel.

## Notes

### Competing Interest Statement

The authors have declared no competing interest.

### Summary of Updates

Corrected author list with correct affiliations.

### Funding

- NIH

