## Supplemental Figures for "Tissue geometry patterns epidermal stem cell fate in murine interfollicular skin"

#### Supplemental Figure S1

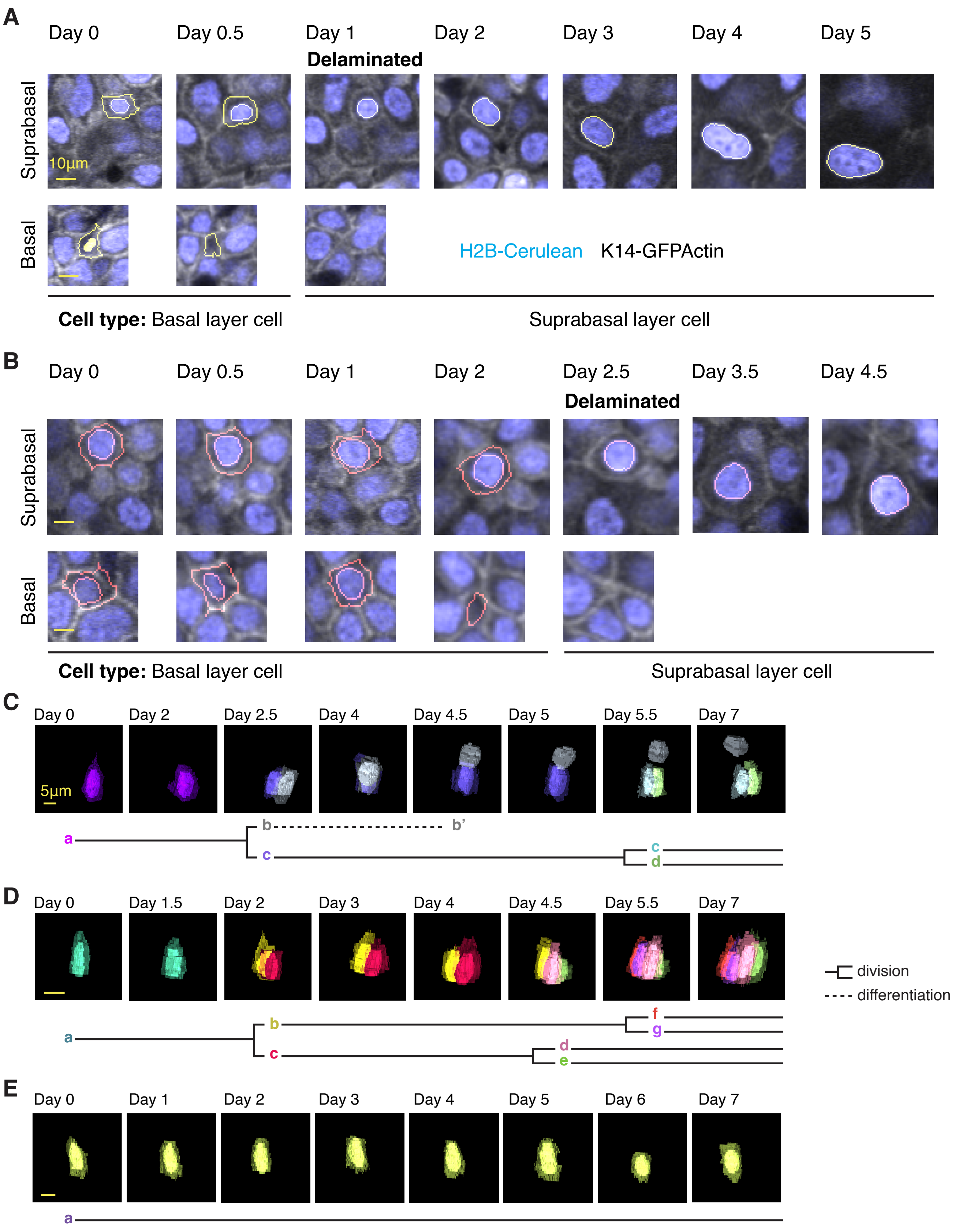

#### Supplemental Figure Legends

**Supplemental Figure S1. Examples of basal layer cell delamination and 4D lineage reconstruction.**

A-B. Two example basal layer cells that delaminated over 7 days imaged in mice expressing H2B-Cerulean and K14-GFP-Actin. The cell and nuclear segmentations of the cells of interest are shown in yellow (A) or orange (B). Suprabasal slices are shown in the top row, while basal slices are shown in the bottom row. The delamination event is manually annotated. Scale bars are 10 μm.

C-E. Three example reconstructed basal layer cell lineages. The top rows show each cell lineage, with all cells within the lineage sharing the same color. For all basal layer cells within the lineage, cell nuclei were rendered as solid surfaces while the cell volumes were rendered as translucent surfaces. For all suprabasal layer cells within the lineage, only the cell nuclei were rendered as solid surfaces. The lineage in (E) is the only cell in the data set that neither divided nor differentiated over 7 days.

#### Supplemental Figure S2

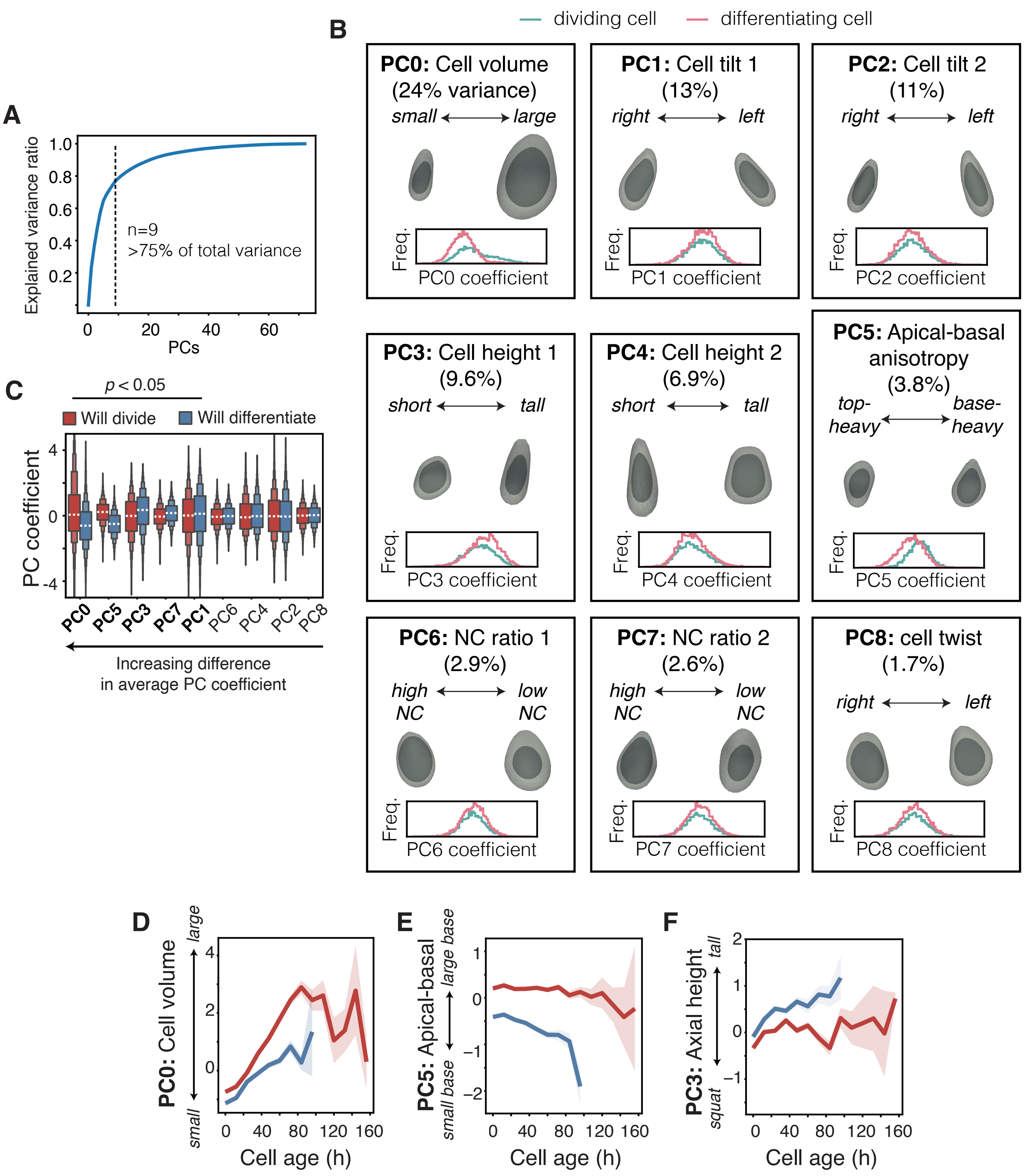

**Supplemental Figure S2. PCA analysis of basal layer cell morphology.**

A. The cumulative explained variance versus the number of morphological PCs included is shown. A cutoff of 9 components was chosen for the analysis, corresponding to over 75% of total variance.

B. Details of each PC are shown. The reconstructed morphology of the principal variation along each PC was reconstructed. Cell volumes are shown in light gray, and nuclear volumes are shown in dark gray. Each PC was given an interpretable name by examining these reconstructed morphologies. In the bottom for each panel, the distribution of PC coefficients for dividing (blue) and differentiating (pink) cells are shown.

C. Distribution of PC coefficients for dividing and differentiating cells. From left to right, these distributions are more different, so they better distinguish dividing or differentiating cells. The distributions are the same as those shown in B.

D-F. The average time series of PC values for PC0, PC5, and PC3, are shown for dividing and differentiating cells.

#### Supplemental Figure S3

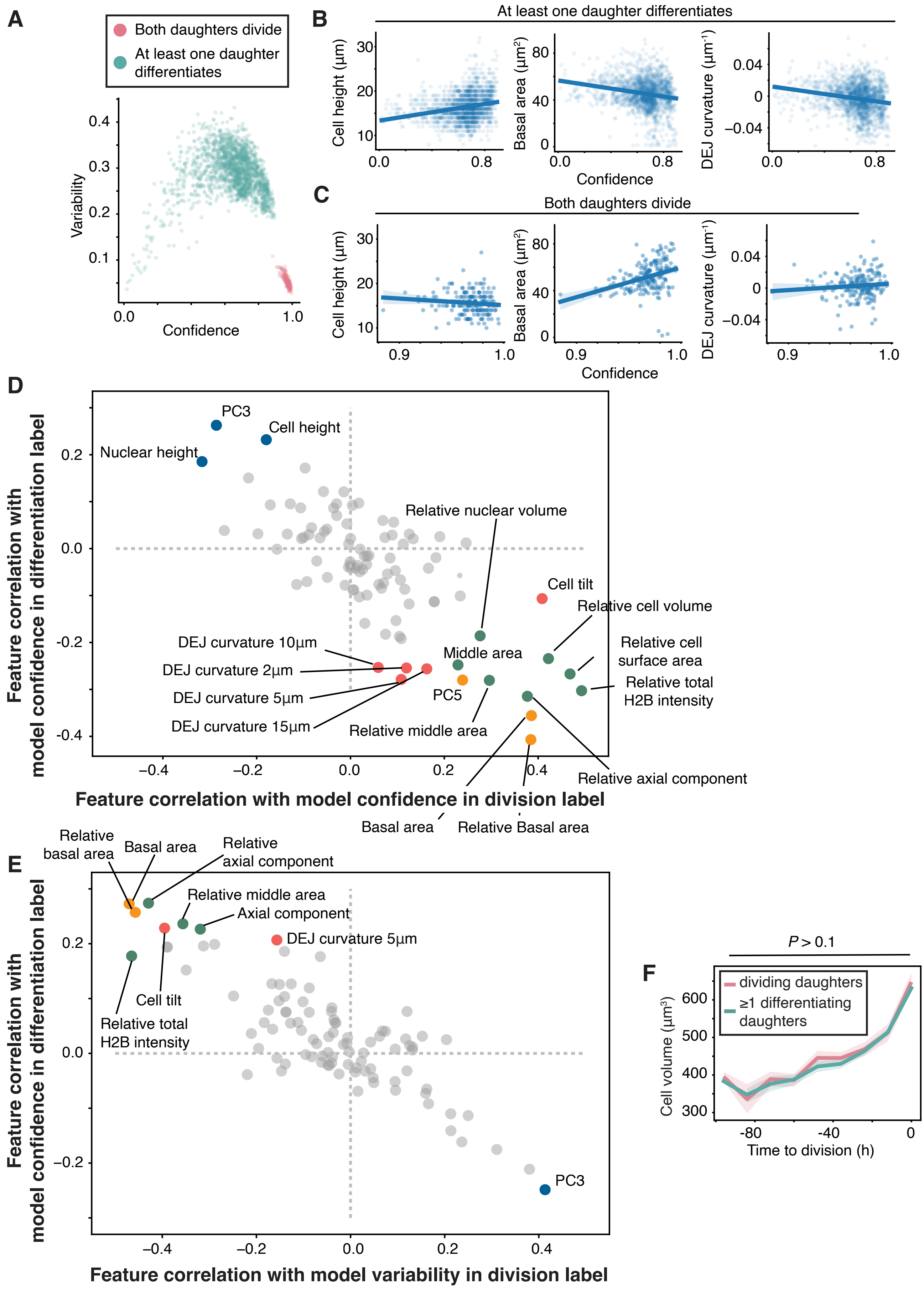

**Supplemental Figure S3. Annotatability analysis of mother cell features that predict daughter cell fates.**

A. The average model confidence and variability as defined by Annotatability (see **Methods**) were quantified across multiple mother cell time points and across multiple independent runs of the model. The model confidence in each cell is colored by the fate of that cell.

B-C. The feature value and Annotatability model confidence are shown for mother cells whose daughters differentiate (B) or divide (C). Solid line denotes a linear regression and the shaded area denotes the 95% confidence interval.

D. The correlation between feature value and model confidence are shown for all input features. This plot is the same as Figure 2E with additional feature name labels. Gray dots are features below a discovery cutoff of $P=10^{-5}$ using Spearman's rho-test.

E. The correlation between feature value and model variability. Gray dots are features below a discovery cutoff of $P=10^{-5}$ using Spearman's rho-test.

F. Cell volume dynamics prior to cell division for mother cells whose daughters either both divide or at least one differentiates. The p-values are calculated using two-sided unpaired Student's *t*-test.

### Supplemental Figure S4

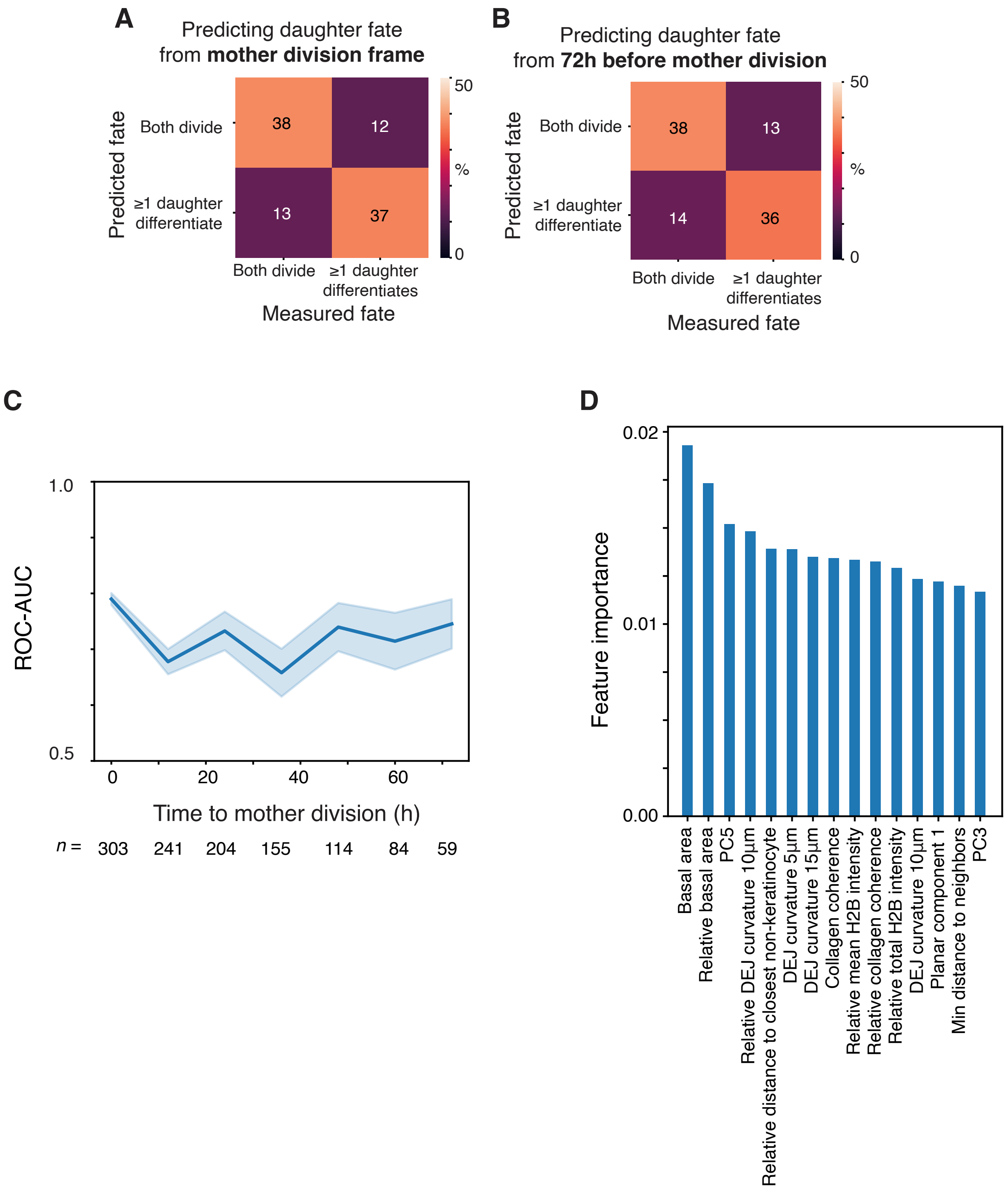

**Figure S4. Random forest classifier analysis of mother cell features that predict daughter cell fates.**

A-B. Confusion matrix for predicting daughter fates from mother features immediately prior to mother division (A), or mother features 72h prior to mother division (B), as evaluated by 80%-20% train test split.

C. The area under the ROC curve for predicting daughter fates from mother features at different time points along the cell history, as evaluated by 80%-20% train test split. Error bars are mean and standard deviation from 1000 independent fits.

D. The average impurity importance of the top 15 features is shown for the random forest classifier model

### Supplemental Figure S5

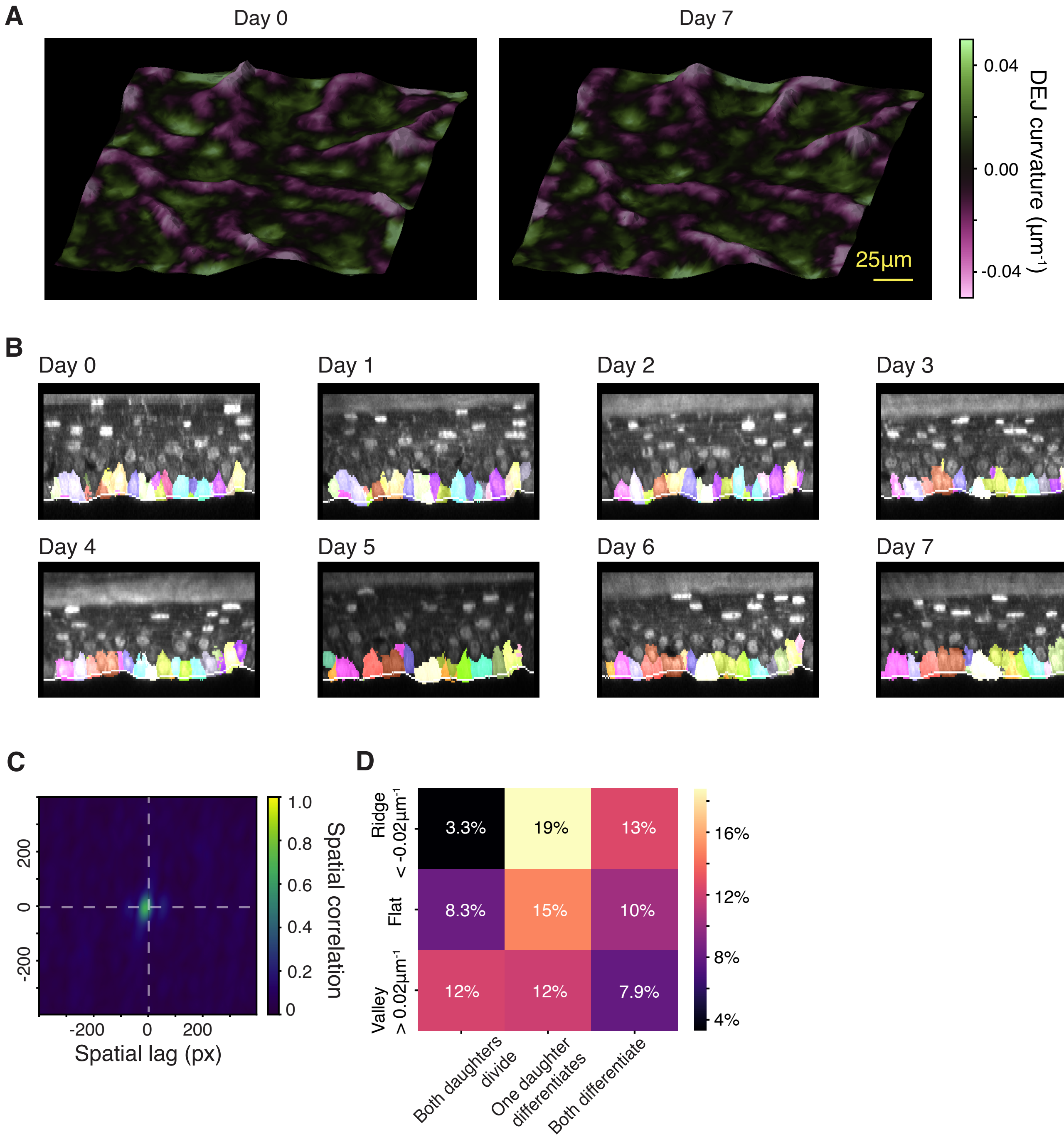

**Figure S5. DEJ curvature persistence and basal layer cell fate.**

A. DEJ surfaces reconstructed from the same hindpaw region measured 7 days apart are shown.

B. A single orthogonal slice from the same image region as (A) is shown from day 0 to day 7. Gray intensity channel is GFP-actin. Individually colored shapes are orthogonal slices of 3D basal layer cell segmentation. White line is the location of the reconstructed DEJ.

C. The spatial correlation function between the DEJ curvatures at Day 0 and Day 18 from **Figure 3D** is shown.

D. The joint probability distribution matrix between daughter cell fate and binned DEJ curvature is shown. The probabilities were calculated from the final timepoint of a mother cell prior to division.

#### Supplemental Figure S6

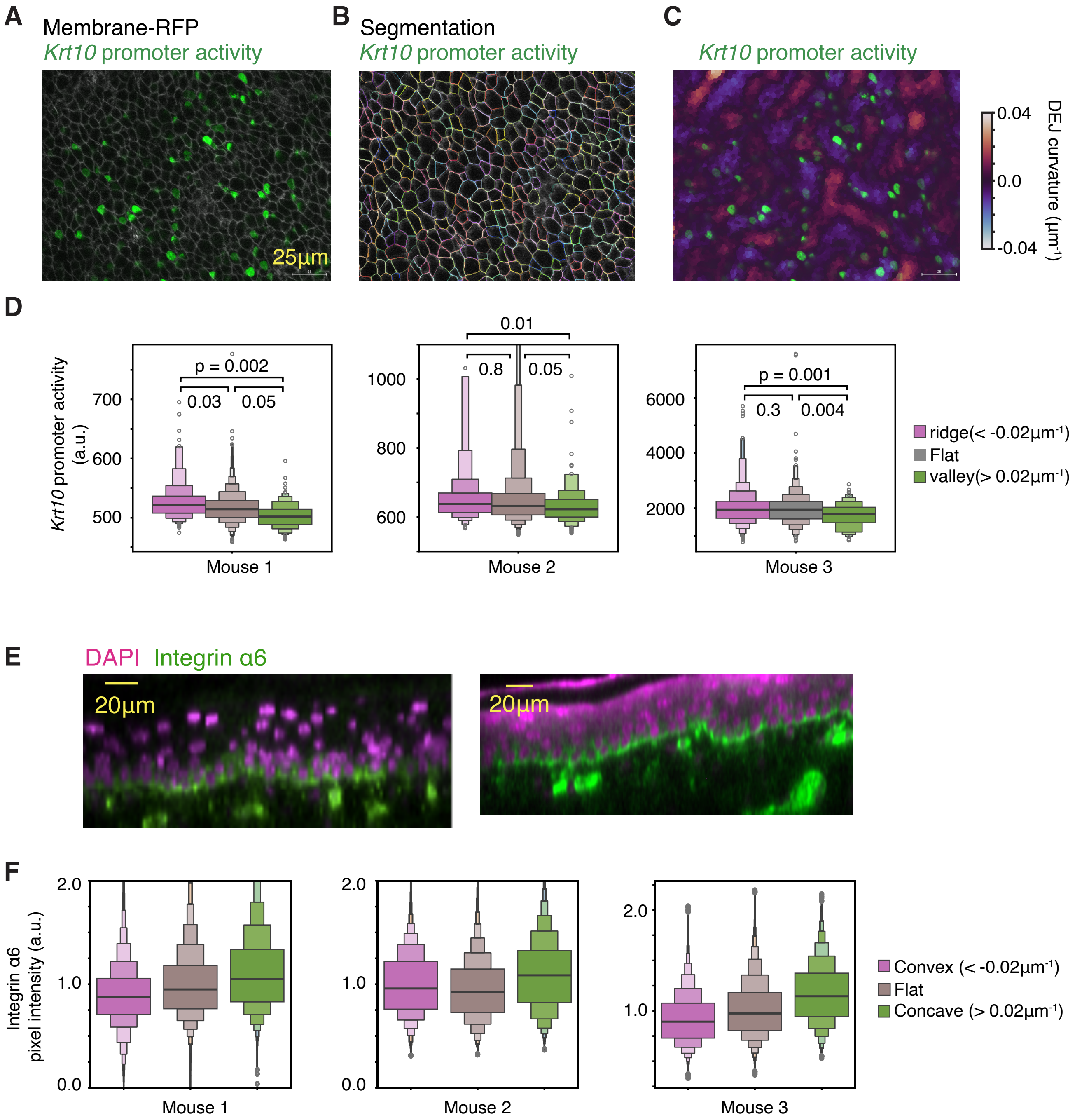

**Figure S6. *Krt10* promoter activity and Itga6 protein expression.**

A-C. Example flattened image of membrane-RFP (A), *Krt10* promoter activity signal (B), and DEJ curvature (C).

D. *Krt10* promoter activity signal intensity is quantified against binned DEJ curvature. The p-values are calculated using two-sided unpaired Student's *t*-test.

E. Orthogonal images of whole mount full-thickness skin stained against Itga6 and DAPI.

F. Itga6 signal intensity is quantified against binned DEJ curvature. The statistical test was not done on a per-pixel basis but on the average pixel intensity per mouse (**Figure 4I**).

#### Supplemental Figure S7

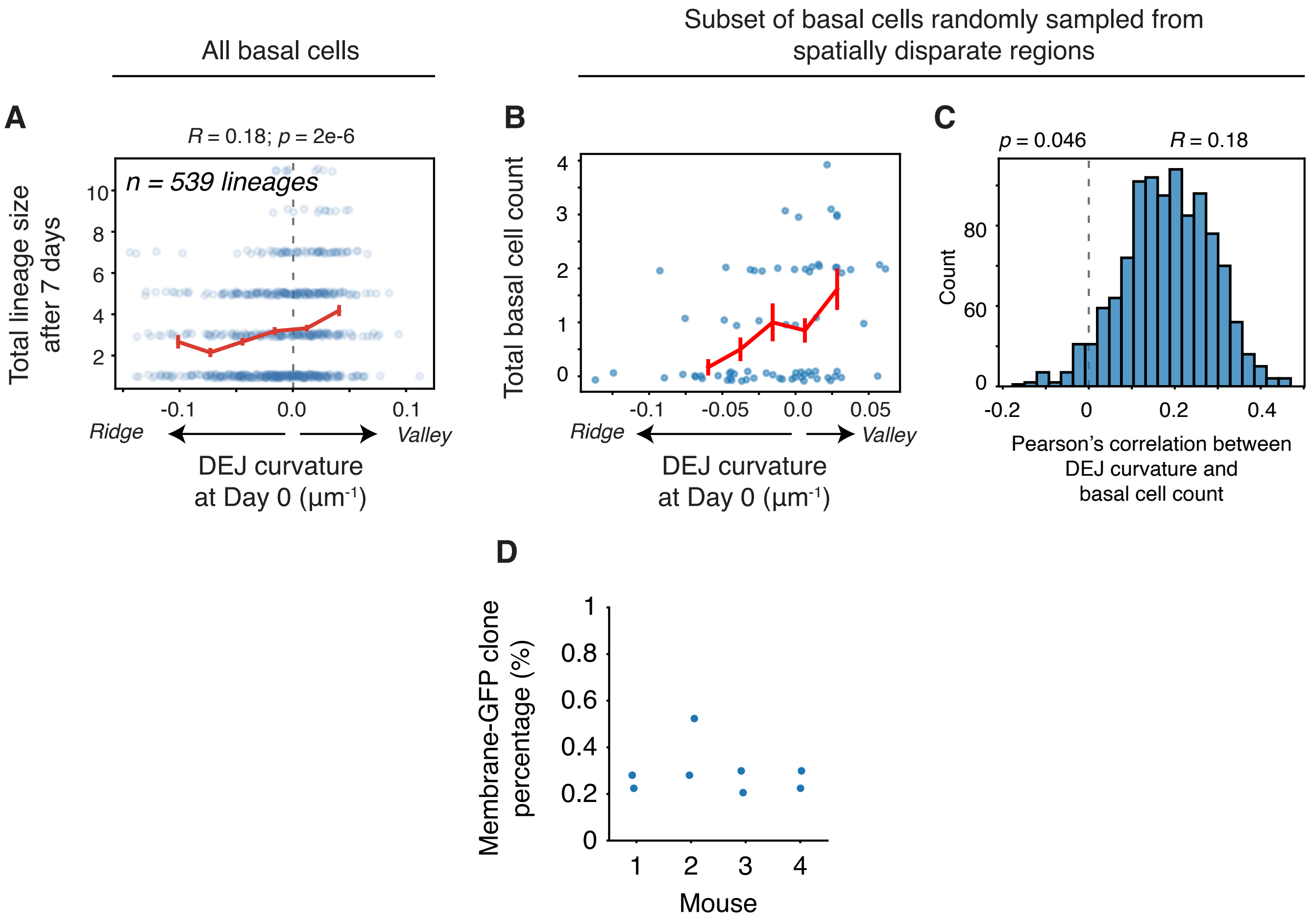

**Figure S7. DEJ curvature and clonal analysis.**

A. Using the densely tracked *in vivo* longitudinal imaging dataset, the total number of descendants after 7 days was quantified against the initial DEJ curvature around a basal layer cell.

B. The imaging field of view at day 0 was blocked into 20 µm x 20 µm regions and a single basal layer cell was chosen at random. The correlation of its initial DEJ curvature and its number of basal descendants is shown.

C. The distribution of Pearson's R between initial DEJ curvature and number of basal descendants is shown for 1000 iterations of randomly spatially blocked lineages. One example is shown in (B).

D. The overall membrane-GFP clone prevalence frequency is shown for 4 different *mTmG* mice.
